# Maternal exposure to a mitochondrial toxicant results in life-long alterations in DNA methylation and gene expression in the offspring

**DOI:** 10.1101/758474

**Authors:** Oswaldo A. Lozoya, Fuhua Xu, Dagoberto Grenet, Tianyuan Wang, Sara A. Grimm, Veronica G. Godfrey, Suramya Waidyanatha, Richard P. Woychik, Janine H. Santos

## Abstract

Mitochondrial-driven alterations of the epigenome have been reported but whether they are relevant at the organismal level remain unknown. The viable yellow agouti mouse (*A^vy^*) is a powerful epigenetic biosensor model that reports on the DNA methylation status of the *A^vy^* locus through the coat color of the animals. Here we show that maternal exposure to rotenone, a potent mitochondrial complex I inhibitor, changes the DNA methylation status of the *A^vy^* locus and broadly affects the liver DNA methylome of the offspring. These effects were accompanied by altered gene expression programs that persisted throughout life. Mitochondrial dysfunction was present in the mothers but not in the offspring until 12 months of age, when electron transport and antioxidant defenses were impaired. These results highlight a putative novel role for mitochondria in nuclear epigenetic remodeling during development, raising fundamental questions about the long-term impact of mitochondrial dysfunction to health and disease.

---

Mitochondrial function affects many areas of cell biology but its impact on epigenetics has just lately gained attention. A decade ago, changes in DNA methylation as a function of mitochondrial DNA (mtDNA) content were first reported (1), and only recently has mitochondrial dysfunction resulting from progressive mtDNA depletion or complex III mutations been shown to alter the histone methylation and acetylation landscapes (2–4). In this context, it was recently shown that the DNA methylation and histone acetylation changes associated with mitochondrial dysfunction influenced gene expression programs. Importantly, these epigenetic and transcriptional effects could be reversed by modulating the tricarboxylic acid (TCA) cycle through genetic or pharmacological means (3, 4). Activation of the methionine salvage pathway was involved in DNA methylation changes while mitochondrial-derived acetyl-CoA was found to influence the histone acetylation landscape (3, 4). Other links between mitochondrial metabolism and the epigenome involving increased reactive oxygen species (ROS) or altered abundance of other TCA cycle metabolites such as α-ketoglutarate or 2-hydroxyglutarate have also been proposed (2, 5). Still, the physiological relevance of the effects of mitochondrial dysfunction on epigenetics *in vivo* remains ill-defined.

The viable yellow agouti mouse (*A^vy^*) is an epigenetic biosensor model that has been widely used in the field of environmental epigenetics. One of its advantages is that the DNA methylation status of the *A^vy^* locus influences the coat color of the animals, which can be used as a visual screen to evaluate DNA methylation *in vivo* (6). Furthermore, because the epigenetic state of the *A^vy^* locus is established before the 3-germ layer separation (7–9), the DNA methylation status of *A^vy^* in the skin reflects the methylation state of the locus in other tissues as well (9). Most notably, work on this animal model has shown that maternal exposure to different diets, physical or chemical agents such as bisphenol A or genistein alter the DNA methylation state of the *A^vy^* locus in the offspring from the exposed mothers, impacting health outcomes in adulthood such as increasing cancer risk or modulating obesity (7, 8, 10). More recently, broader epigenetic analysis revealed that histone acetylation is also modulated in the *A^vy^* locus (11).

The *A^vy^* animals carry a germline insertion of an intracisternal A particle (IAP) upstream of the coding exons of the *agouti* gene. Activation of a cryptic promoter in the IAP hijacks the regulation of agouti expression, resulting in high level transcription of a fusion IAP-agouti message that produces the normal agouti protein throughout the animal (9). The ectopic expression of this fusion transcript results in animals with a yellow coat color, obesity and type II diabetes (9, 12). DNA methylation at the long terminal repeats (LTR) of the IAP effectively shuts down the ectopic expression from the cryptic promoter, which results in normal expression of *agouti* in the skin. Animals with a methylated IAP LTR exhibit pseudoagouti coat and are indistinguishable from wild-type animals. Cellular mosaicism within a single animal can also arise, leading to different degrees of mottling of the coat in animals that carry the *A^vy^* allele and intermediate phenotypes (9).

The mechanism underlying the epigenetic changes that lead to the variable penetrance of the *A^vy^* allele remains unknown but cannot be attributed to variability in the genetic background of individual animals since experiments were conducted on an inbred background. It is also unclear how the physical and chemical treatments of the mothers during pregnancy lead to changes in the epigenetic status of the *A^vy^* allele in the offspring. Here, we tested the hypothesis that subtle modulation in mitochondrial function during development underlies the changes in the DNA methylation status of the entire nuclear genome, including the *A^vy^* locus. By exposing *A^vy^* animals perinatally to rotenone, a pesticide that specifically targets the site of NADH oxidation on mitochondrial complex I, we identified changes in the DNA methylation of the *A^vy^* locus in the skin and in the total methylome in the liver. More notably, perinatal rotenone treatment impacted DNA methylation in thousands of loci in the liver, including those that should normally change as the animals age. We also documented the significant remodeling of the liver transcriptome, throughout life, that result from rotenone treatment.

## Results

### Maternal exposure to rotenone leads to hypomethylation of the *A^vy^* locus in the offspring

We started with dose-response experiments in a 2-week exposure protocol to define the concentrations of rotenone that would inhibit complex I but would not exhibit toxicity to C57BL/J animals. The National Toxicology Program (NTP) previously reported that a 2-year exposure of B6C3F1 mice to 600 (120 mg/Kg) or 1,200 ppm (240 mg/Kg) rotenone through the diet was not toxic, with only minor effects on body weight and, unexpectedly, decreased frequency of liver cancer in male mice (13). Thus, our initial exposures involved 10, 50, 150, 300, and 600 ppm rotenone. Prior to studies, feasibility of formulating rotenone at these concentrations in the feed and its stability over the time-frame required for the experiments were established (see Methods). Unlike the B6C3F1 animals used by the NTP for the 2-year cancer bioassay, rotenone feeding at 600 ppm was highly toxic to C57B6/J mice. Animals showed signs of frailty, hair loss and decreased food consumption within 2 days of exposure (Fig. S1A). Multiorgan toxicity was observed after 2 weeks of feeding even with 300 ppm (see report in Appendix), which led to the removal of these concentrations from the study. Based on the lack of toxicity and the degree of complex I inhibition in isolated liver mitochondria from exposed females (Fig. S1B), we used 10 and 150 ppm rotenone, which based on the daily food consumption of this strain of mice is estimated to be equivalent to 2mg/Kg and 30 mg/Kg, respectively. Final number of animals utilized was based on power analysis.

A diagram of the experimental design is shown in Fig. 1. Briefly, 180 (8-week-old) dams were exposed to control- or rotenone-supplemented diets for 8 full weeks starting 2 weeks prior to mating, during the 3 weeks of gestation and throughout the 3 weeks of lactation; males carrying the *A^vy^* mutation were mated with the rotenone-treated or control females and were only exposed during this period. Offspring from the females were phenotyped for their coat color and put on the control diet at weaning (post-natal day 21, PND21). The offspring were followed and analyzed at specific time intervals over the course of approximately two years. A total of 669 offspring were generated, about half of which carried the *A^vy^* mutation. Table 1 summarizes information from the control- or rotenone-fed offspring. While we did not find overt toxicity in the 150-ppm feeding, this dose of rotenone altered most parameters evaluated (Fisher’s exact test: p<0.0001), including pregnancy rate, litter size and pup survival rate relative to the control group. Control and 10 ppm-cohort were essentially indistinguishable although rotenone treatment unexpectedly increased survival rate (p<0.0001; Table 1).

**Figure 1.**
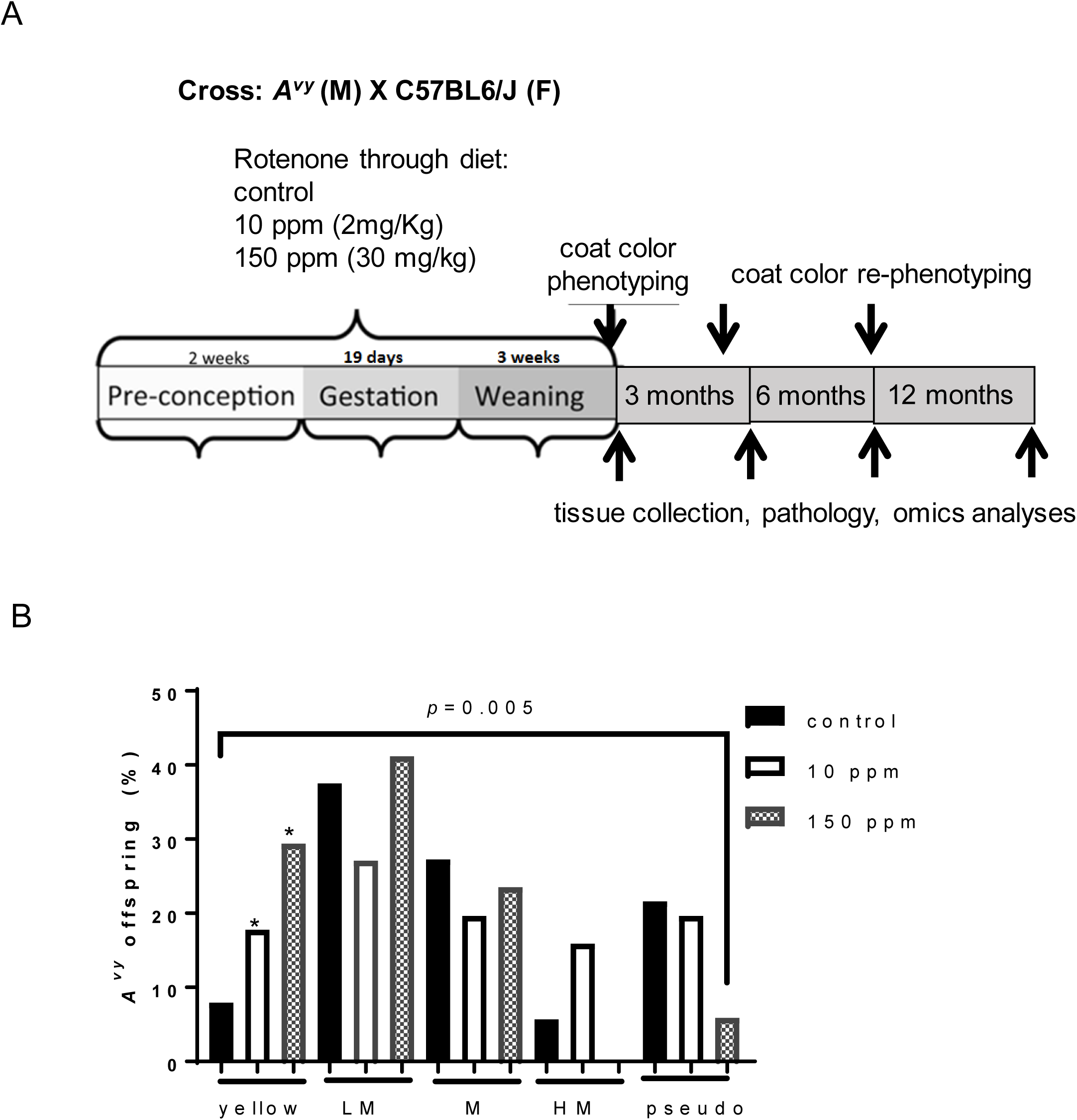
Maternal rotenone exposure alters the frequency of yellow *A^vy^* offspring. **(A)** Rotenone was administered in two different doses through the diet (AIN-93G) to both females and males; females started exposure 2 weeks prior to conception until weaning while males were exposed only while mating. Pups were phenotyped for coat color based on visual inspection at weaning, 5 and 12 months. Tissues were collected for various types of analyses throughout the course of the experiments. (B) The frequency of animals in each of the 5 categories of the *A^vy^* coat color phenotype was calculated based on the number of animals in that category relative to the total number of alive mutant animals in the cohort in each experimental group (N=212). Y – yellow, LM – lightly mottled, M – mottled, HM – heavily mottled; pseudo – pseudoagouti. Chi-square test was applied to determine significance of distribution differences within each category across doses relative to the control; significance is shown with * where p=0.023. Somer’s test was applied to determine significance in the order of increasing methylation as a function of the category i.e., methylation Y<LM<M<HM<pseudo based on dose of exposure. Only the highest dose showed significance; p=0.4 for 10 ppm and p=0.005 for 150 ppm.

**Table 1:** Summary of endpoints in the control or rotenone-treated cohorts.

| endpoint | control | 10 ppm | 150 ppm |
| --- | --- | --- | --- |
| pregnancy rate | 48/60 (80%) | 51/60 (85%) | 43/60 (71.7%)* |
| days to delivery | 25±4.8 | 25.8±6.3 | 29±9* |
| number of litters | 40 | 39 | 28* |
| litter size | 6.96±1.3 | 6.92±1.6 | 4.3±1.6* |
| number of pups | 278 | 270 | 121* |
| pup survival rate | 76.50% | 84.30%* | 50.05%* |
| a/a offspring | 47.62% | 51.36% | 41% |
| male offspring | 48.81% | 47.27% | 66% |
| weight at weaning | 8.37±1.7 | 7.58±1.08 | 5.65±0.44* |
| alopecia | 0 | 1 (0.3%) | 6 (4.96%) |

Visual inspection of the *A^vy^* offspring revealed a dose-dependent increase in the frequency of fully yellow-coated animals when females and males were pooled (Fig. 1B) or scored separately (Fig. S2A), which is indicative of a change in DNA methylation at the *A^vy^* locus. Because coat color can be arranged in order of increasing methylation from yellow, to lightly mottled, to mottled, to highly mottled and to pseudoagouti, we applied statistics to test whether the percentages in this order correlated with the dose. Somers’ test showed a significant correlation (p=0.005) for the 150 ppm but not for the 10 ppm (Fig. 1B). We found that the coat color of individual animals was stable as the animals aged, which suggested that the underlying DNA methylation status of the *A^vy^* locus in the hair follicle was stable over the 2-year period of observation. This is consistent with previous reports using this same mouse model (14). As previously shown (9, 12, 15), animals with varying degrees of yellow color became obese within 8 weeks.

These data provide unequivocal evidence that developmental exposure to rotenone impacts the methylation status of the *A^vy^* locus in a way that affects the coat color distribution in the offspring of the treated female mice.

### Maternal mitochondrial complex I inhibition alters genome-wide DNA methylation dynamics in the liver over the lifespan of the offspring

Having determined the effects of rotenone exposure on the DNA methylation of the *A^vy^* locus in the pups exposed *in utero* and throughout lactation, we next set out to define if the epigenetic status of other loci were also affected in the offspring. To this end, we performed whole genome bisulfite sequencing (WGBS, 7X genome coverage per animal) at a single nucleotide resolution in the livers of the treated offspring. For this analysis, we only used black animals and the offspring exposed to 10 ppm rotenone to avoid confounding effects associated with the metabolic syndrome of yellow animals and the potential toxicities associated with the 150-ppm exposure (Table 1). After sequencing the DNA from males and females at PND22 and at 6 months, we found no genome-wide gender-specific differences in DNA methylation (Fig. S2B). Thus, further analyses involved only females.

At least 3 females were sequenced at PD22, 6, 12 and 18 months. The impact of perinatal rotenone exposure to genome-wide DNA methylation was assessed based on the identification of differentially methylated regions (DMRs) between the two groups (see Methods). Each line on the heatmaps (Fig. 2) displays DMRs that were significantly different at any given time in each group; the same locus was displayed over the course of the experiment to visualize the dynamics of the aging DNA methylome. The data was clustered based on the patterns followed by the DMRs over the time-frame of the experiments in each group separately. Under our experimental conditions, DMRs represented less than 0.5% of the mouse liver autosomal DNA methylome (Fig. S3A), which is consistent with recent findings obtained in different strains of mice (16).

**Figure 2.**
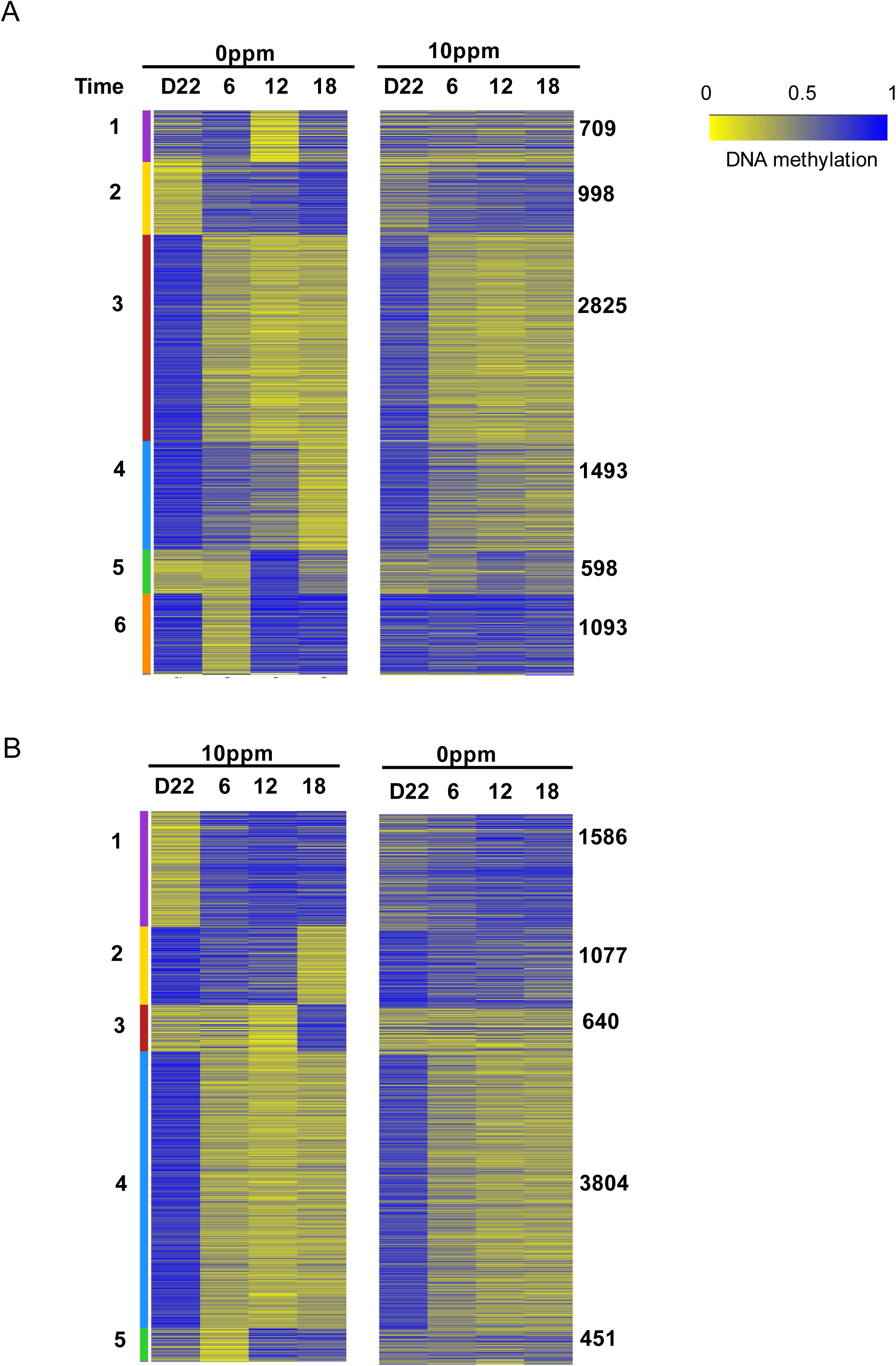
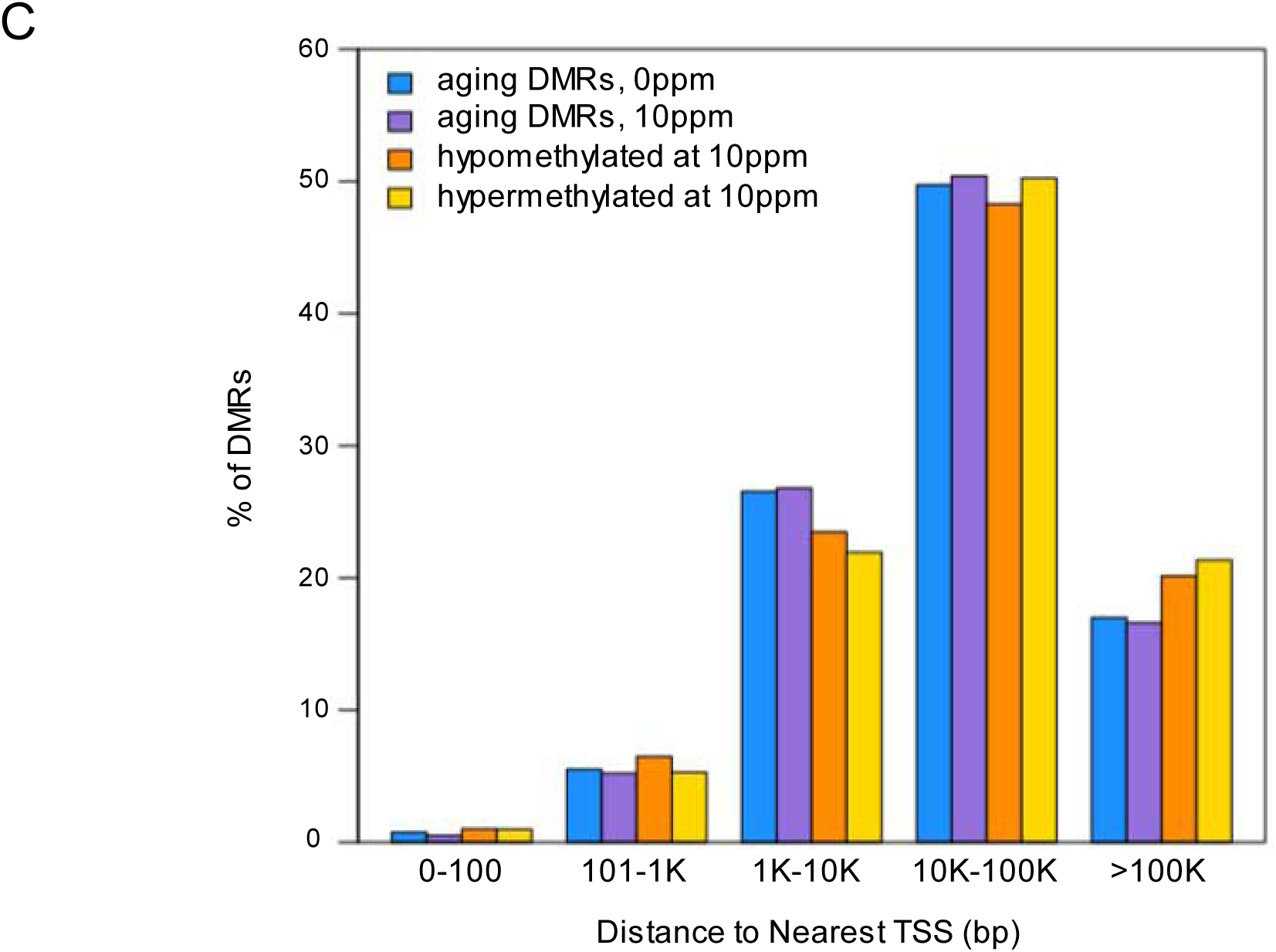
Perinatal rotenone exposure modulates the normal changes of the aging liver DNA methylome. Whole genome bisulfite sequencing (WGBS) was performed in the livers of females (N=3/group/time). Heat map displays differentially methylated regions (DMRs). Blue: hypermethylated, yellow; hypomethylated. Color bars on left represent clusters (numbered on left); the numbers on the right of each cluster indicate the number of DMRs that follow that pattern. On top of panels time is shown, 6, 12 and 18 refer to months. D22 = post-natal day 22; (A) data based on control animals, (B) results are based on DMRs identified in the perinatally-exposed rotenone cohort. (C) Distance of identified DMRs to transcriptional start sites (TSS) based on Gencode VM18.

We found 7,716 DMRs when analyzing the DNA methylome of the control group (Fig. 2A, left panel). Herein we call these ‘aging DMRs’ since they reflect changes in the methylome as a function of age. Some loci started hypomethylated at the early time point(s) but became hypermethylated over time (cluster 2) while others followed the opposite trend (e.g. clusters 3 and 4). Other DMRs showed intermediate patterns (see clusters 1, 5 and 6), highlighting that the aging liver DNA methylome is highly dynamic. Most of these same genomic regions were not altered in the rotenone-exposed animals, with only DMRs following the pattern of cluster 3 showing similar changes (Fig. 2A, right panel). Analysis of the rotenone-exposed offspring showed that the number of DMRs was similar and involved 7,558 loci (Fig. 2B, left panel). These rotenone-affected DMRs clustered in 5 patterns instead of 6 as was observed in the control group (Fig. 2B). When evaluating the DNA methylation status of these same 7,558 loci in the controls, it was clear that other than DMRs following the pattern of cluster 4, no significant changes were observed in the other regions (Fig. 2B, right panel).

Collectively, these results show that perinatal rotenone exposure significantly impacted the normal dynamics of DNA methylation that occurs as the liver ages.

### DNA methylation changes occur in loci that are relevant to liver physiology

We next assigned genes to the DMRs to define biological processes potentially affected by aging and the rotenone treatment. We arbitrarily assigned a DMR to the closest transcriptional start site (TSS) given that most DMRs were between 1-100Kb of TSSs (Fig. 2C). A full list of the DMRs, their coordinates and closest gene and TSS and whether they were hyper- or hypomethylated after rotenone exposure are presented in Table S1. The degree of DMR overlap between the two groups as well as the overlap in the number of DMR-assigned genes can be found in Figs. S3B and C. We performed pathway enrichment analysis using KEGG (Kyoto Encyclopedia of Genes and Genomes) and the aging DMRs (Fig. 2A, clusters 1-6) or the DMRs identified in the rotenone-cohort (Fig. 2B, clusters 1-5). We found that most pathways while common between the two groups ranked differently (Table S2). For instance, the first enriched pathway when using aging DMRs was cholesterol metabolism followed by bile secretion; these were ranked 29 and 6, respectively, when considering animals that had been through rotenone exposure (Table S2). Conversely, drug metabolism was ranked 1 in the rotenone-exposed cohort while hippo signaling, which regulates organ size through cell proliferation and apoptosis was ranked 3^rd^. In the controls, hippo signaling was ranked 69 while drug metabolism was ranked 18^th^ (Table S2). Because ranking is a function of how many genes within the pathway are represented in the dataset, these results are consistent with the different prioritization and level of engagement of biological processes taking place in the liver in the non-exposed or rotenone-exposed animals.

DMRs can impact transcription factor (TF) binding. Likewise, genomic DNA sequences within a group of select TF bindings sites can influence the local DNA methylation status (16). Hence, we next applied a motif analysis algorithm (HOMER) to determine which TFs were enriched at the identified DMRs. Irrespective of whether the DMR was hyper- or hypo-methylated in the rotenone-exposed cohort, we found significant enrichment for several TFs, including some associated with the regulation of mitochondrial function, e.g. NRF-1 (Nuclear Respiratory Factor 1). Other enriched TFs included ARNT (Aryl Hydrocarbon Receptor Nuclear Translocator), a protein that binds to ligand-bound aryl hydrocarbon receptor and aids in its nuclear localization to promote the expression of genes involved in xenobiotic metabolism, EGR1 (Early Growth Response Element 1) that is involved in differentiation and mitogenesis, and HNF6 (Hepatocyte Nuclear Factor 6), which is a liver-specific TF (Fig. S3D).

Taken together, these results show that DMRs enrich for pathways and TFs that are not only relevant to liver biology but also to mitochondrial function and xenobiotic metabolism, which is in line with exposure to a mitochondrial toxicant.

### Developmental rotenone exposure alters the normal age-related liver gene expression program prior to signs of pathological changes

We next examined the effects of maternal rotenone exposure on the transcriptome of the liver of the offspring. To this end, we performed RNA-seq on the same samples utilized for WGBS; PND22 was used as the reference for time-point comparisons within each treatment group and between rotenone-treated and the controls. We started by defining the effects of rotenone at baseline (i.e., PND22), which was when rotenone exposure was terminated. At this time 252 genes were differentially expressed (DEGs), some of which involved mitochondrial function (Table S3 and Fig. 3A). Although rotenone at high doses has been previously shown to affect tubulin polymerization (17), no genes involving tubulin metabolism were detected as differentially expressed at baseline (Table S3). The pathways enriched by the baseline DEGs can be found in Table S2 and are also shown in Fig. 3B (in grey). It was notable that, in addition to signaling and cell cycle regulation, the cholesterol and mevalonate pathways were enriched. Changes in fatty acid metabolism are recognized as effects of chronic rotenone exposure (18). Also, DMRs too enriched for cholesterol metabolism (Table S2).

**Figure 3.**
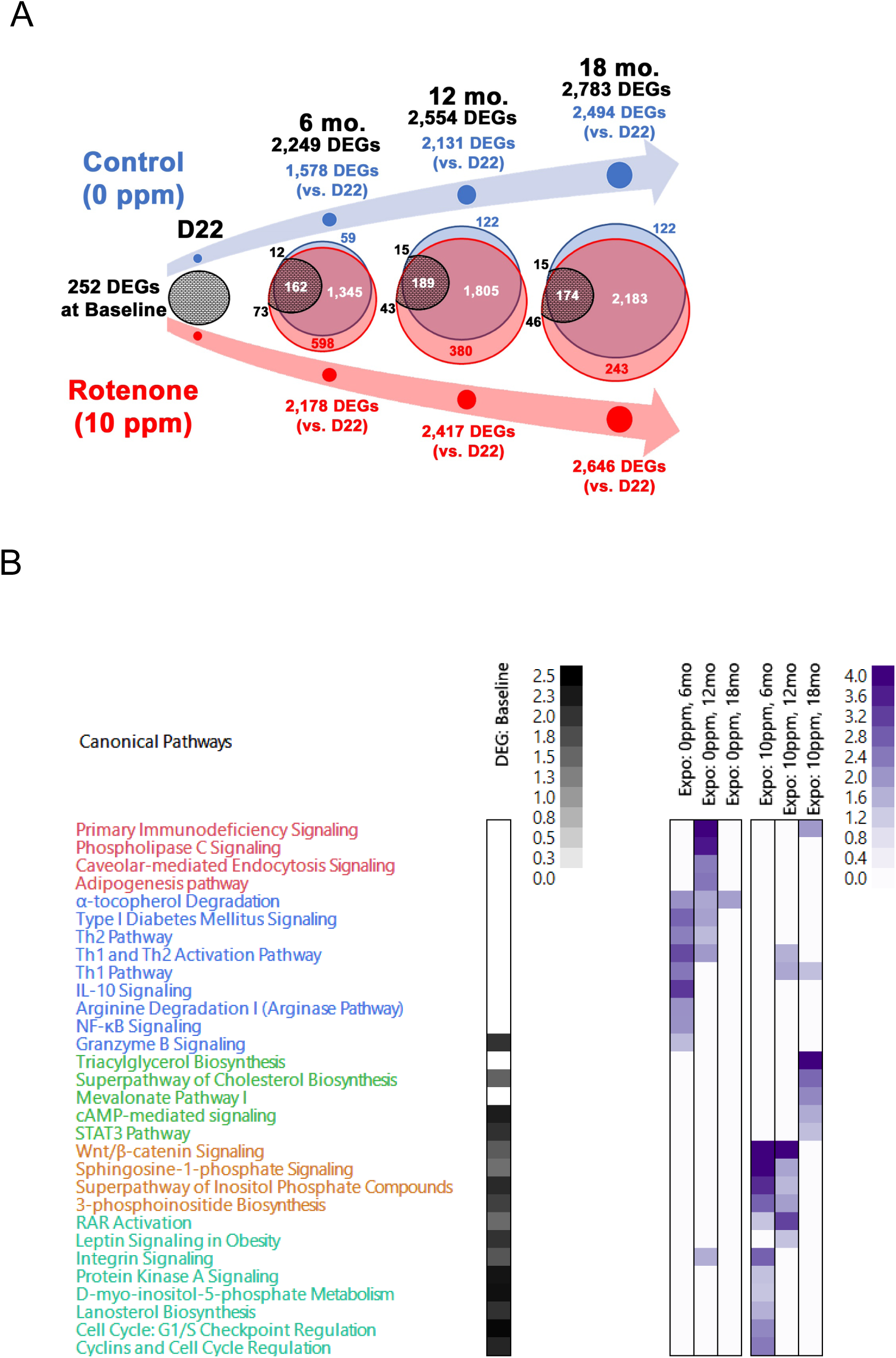
Life-long changes in the liver transcriptome are detected after perinatal rotenone exposure. The transcriptome of the livers submitted to WGBS was evaluated using RNA-seq. N=3/time/group. (A) Venn diagram depicts the overlap of differentially expressed genes (DEGs) identified at each time point in the experimental groups. Black numbers on top depict all DEGs identified. Baseline DEGs were obtained by comparing rotenone to control cohort at post-natal day 22 (D22). Blue depict the number of DEGs identified in the control cohort and in red those identified in the rotenone-fed counterparts. Data at each time point are relative to D22 within each group. The black circles show the number of DEGs identified at baseline that were still differentially expressed at later times and the amount that was common to both groups; number above or below these circles (in black) depict baseline DEGs unique to control (upper) or rotenone-fed cohort (lower). (B) Ingenuity Pathway Analysis (IPA) was used to define biological processes enriched by the DEGs. Black heatmap was obtained when using baseline DEGs, purple heatmaps were generated using DEGs identified as uniquely expressed in the control group or in the rotenone-exposed cohort over time, respectively.

Most genes, ∼60-87%, were shared between the two groups (Fig. 3A) with similar fold-changes (Table S4), reflecting a highly coordinated gene expression program in the liver during aging. Despite the similarities in the degree of individual gene expression changes at any given time between the two groups, it was notable that many DEGs were already changed at baseline in the rotenone-fed cohort. Most interesting, all but two genes (Slc4a1 and Rnf133) changed in opposite directions between baseline and the other time-points, which resulted in a large net change in their gene expression. The common genes differentially regulated in both experimental groups were enriched for cell cycle regulation, DNA repair, nucleotide synthesis, hepatic steatosis and various signaling pathways (Table S4).

Some genes were only changed in the control animals (Fig. 3A); these were primarily involved in signaling and in the immune response (Fig. 3B in purple and Table S5). The latter may reflect an increased amount of resident immune cells or their infiltration as a sign of inflammation. Irrespective of the exact reason, this normal response was blunted by perinatal rotenone exposure. Likewise, hundreds of genes were uniquely differentially expressed in the offspring of rotenone-exposed mothers, an effect that was observed at 6, 12 and even 18 months after exposure ceased (Fig. 3A). This unique but altered gene expression program enriched for signaling and metabolic pathways, which were not changed in the progeny of animals fed the control diet but that were modulated in the offspring of exposed animals already at PND22 (Fig. 3B in purple and Table S6).

The effects on the transcriptome prompted us to dissect the livers to interrogate signs of pathology. Given that most transcriptomic changes were observed at 6 and 12-month-old animals, we analyzed animals at the 12-month time-point. One expectation based on the gene expression profiles was that the perinatal rotenone exposure would increase lipid accumulation while decreasing signs of inflammation, at least in the liver. Ten animals per group were examined, and tissue and body weight were recorded at time of necropsy. No changes in body weight or organ/body weight ratio were identified for any of the tissues analyzed, which included the heart, liver, kidneys, pancreas and thymus. No histological changes were found in the brains, consistent with the lack of neuronal toxicity with the dose of 10-ppm. Histopathological assessment of the liver, heart and lungs identified changes in all organs, including immune cell infiltration and fatty acid accumulation. However, no statistical differences were found between the rotenone-treated or control groups (Table S7), which may reflect the sample size.

### Altered DNA methylation patterns correlate with changes in gene expression

Many of the pathways enriched using the DMRs were also found when using the DEGs, suggesting a relationship between the DNA methylation and gene expression changes observed. To better understand whether these genomic changes were occurring in parallel or correlated with each other, we capitalized on the fact that DNA methylation and RNA-seq analyses were performed on the same samples. We started by determining the overlap between DMR-assigned TSS (closest physical TSS) with the coordinates of the TSS of the DEGs. When analyzing all genes, we found that ∼29% of all DEGs corresponded with the nearest TSS to DMRs (Table S8 and Fig. 4A, left). When considering each time-point, the overlap was closer to 8-10% depending on the group (Fig. 4A, right).

**Figure 4.**
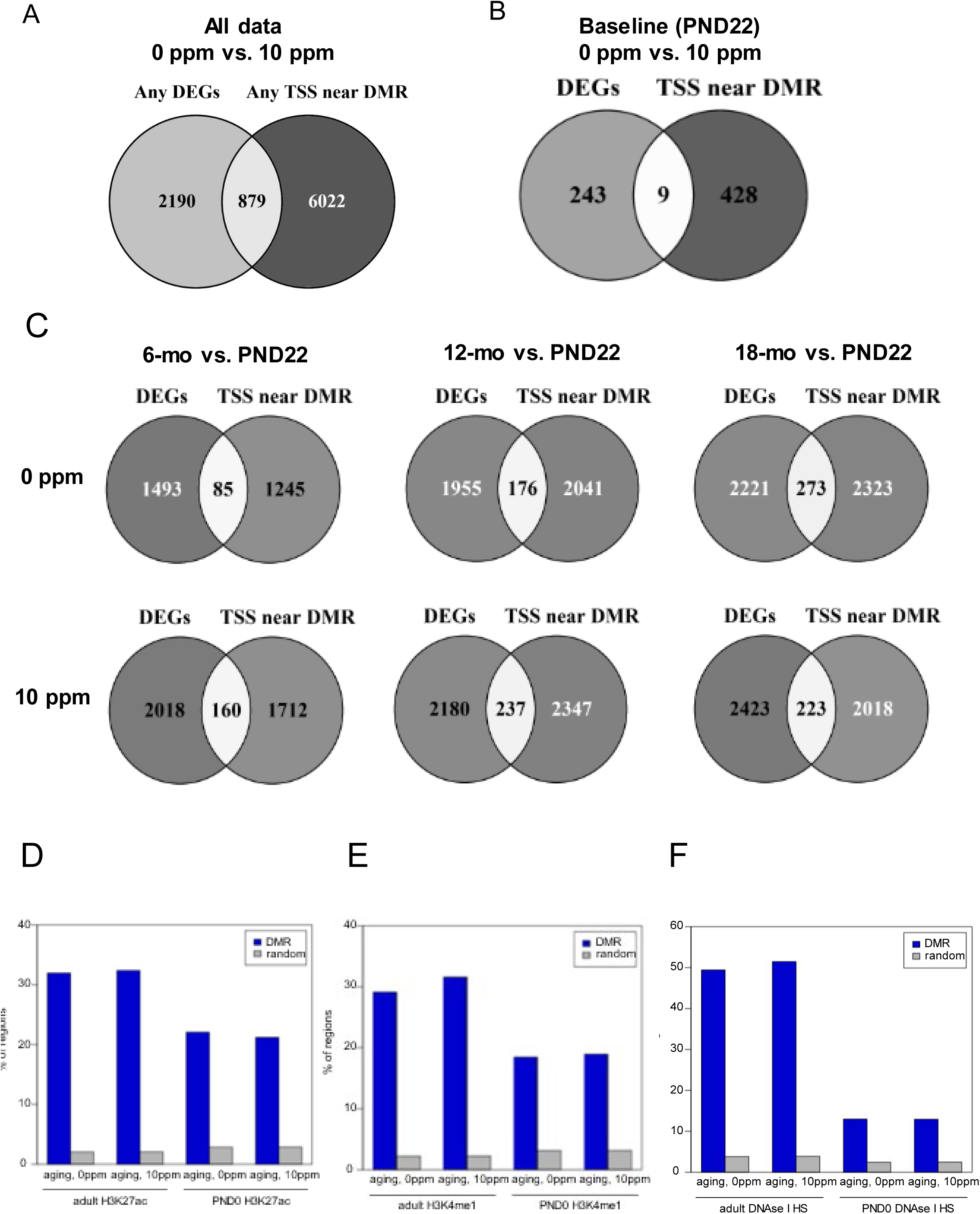
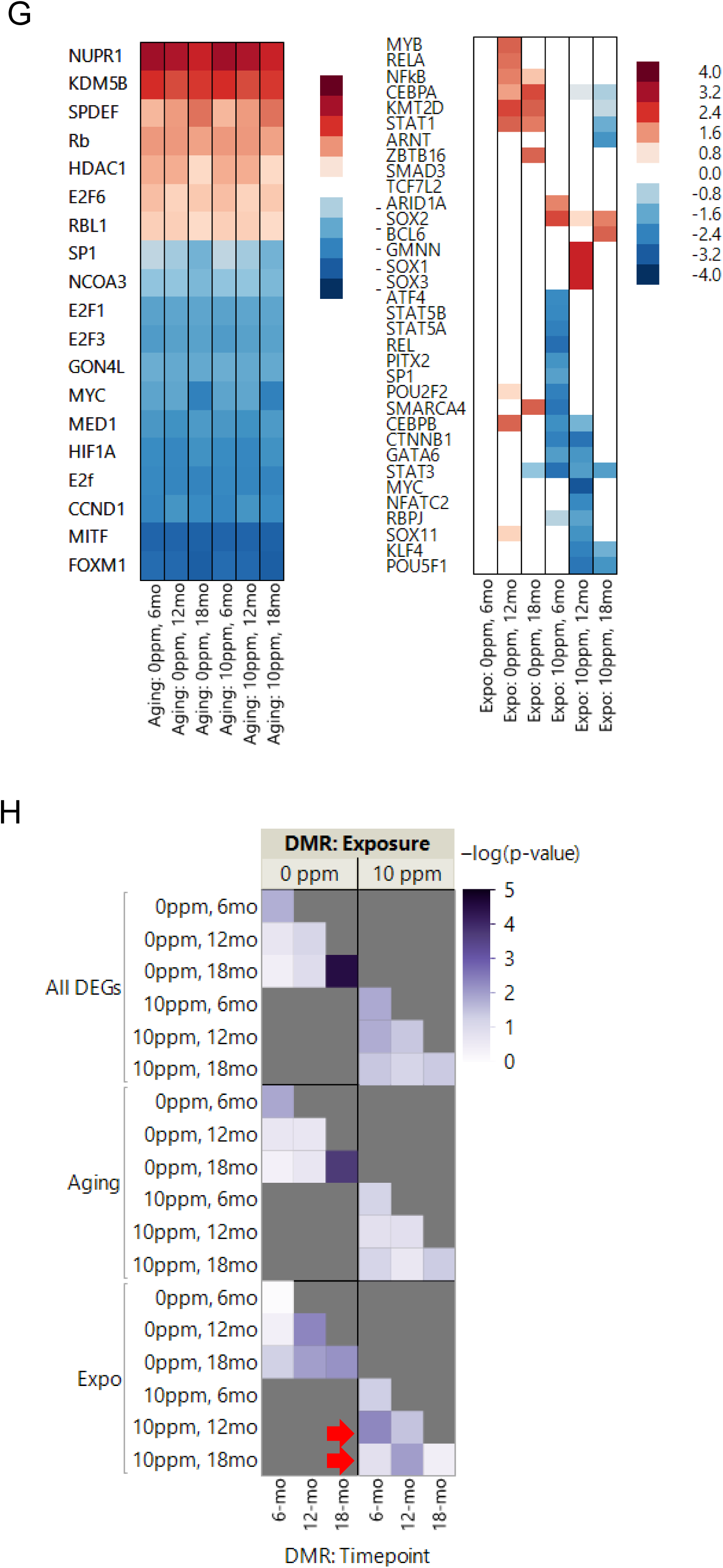
Differentially methylated regions overlap with differentially expressed genes and regulatory DNA. Venn diagram depicts the overlap between DMRs and the DEGs based on closest TSS using all data compiled (A) or per time-point. (B and C). D and E panels show graphs depicting the overlap of DMRs with regions enriched for the enhancer marks H3K27ac or H3K4me1 (ENCODE) or (F) DNase hypersensitive sites (UCSC Genome Browser). These data were compared to the average overlap observed for 1,000 iterations of size-matched random genomic sequences. (G) IPA was used to predict upstream regulators of the gene expression programs based on all genes identified in either the control or rotenone-exposed cohort (left) or based on genes differentially expressed exclusively in the control or in the rotenone-fed offspring (right). (H) Heatmap of significance scores [−log(p)] for enriched representation of genes with DMRs with respect to DEGs. Data were obtained across groups based on time points and involved all genes (upper panels), aging- (middle panels) or rotenone-responsive DEGs (lower panels). Red arrows indicate a left-shift on the timing of DNA methylation relative to when genes were differentially expressed.

Next, we considered the possibility that a DMR may regulate the expression of a distant gene. Recently, DMRs were shown to largely overlap with enhancers (16), which can drive transcription of close or distant promoters (19). Thus, we compared the DMRs with coordinates of enhancers based on their frequent association with the modified histones H3K27ac and H3K4me1 and/or overlap with DNAse hypersensitive sites (20, 21). Using publicly available ENCODE data from liver of C57B6 neonate (PND0) or adult animals (8-10 weeks), we found a significant overlay between all aging or rotenone-affected DMRs and loci enriched for H3K27ac or H3K4me1, reaching ∼30% on average (Fig. 4B and C). Depending on the cluster the DMR belonged to, the overlap was closer to 50% (Table S9). Similarly, ∼50% overlap was found between aging or rotenone-DMRs with DNAse hypersensitive sites (Fig. 4D) with some DMRs from specific clusters reaching almost 80% overlap (Table S9). These values are in accordance with or even exceed those recently identified using liver WGBS data (16), and support the conclusion that the association between DMRs with regulatory genomic DNA regions is not random.

A fundamental step to define whether some of these enhancer/DMRs regulate gene expression would be to define their interaction with the target promoter. HACER, an atlas of human active and *in vivo*-transcribed enhancers was recently created (22). This single data repository integrates results from FATOM5 CAGE profiles, GRO/PRO-seq data, ENCODE ChIP-seq data and chromosome conformation capture technologies with validated interaction from high-throughput experiments (22). As such, it provides high confidence *in silico* information on TF-enhancer binding and enhancer-promoter interactions. We reasoned that if the DMRs we identified were regulating gene expression through enhancers, then TFs that recognize those DMRs/enhancer sites should overlap with the TFs predicted to drive the transcriptional programs identified. To this end, we first used IPA to predict the upstream regulators of the gene expression programs (Fig. 4E) and then utilized HACER to predict TF-enhancer binding.

HACER was developed based on human data (22). Thus, we first obtained the human coordinates corresponding to the mouse DMRs we identified. Then, we queried the DMRs at each time point for enhancer overlap and TF-binding. Consistent with our comparison to the ENCODE data, we found that several DMRs in the control- or rotenone-fed cohorts overlapped with mapped enhancers (Table S10). Most importantly, we found that the TFs identified as bound to these enhancers overlapped with 7 of the top 15 TFs predicted to be upstream regulators of the gene expression program, including MYC, SP1, KDM5B, HDAC1, FOXM1, E2F1 and E2F6 (Fig. 4E). Some of these enhancer-bound TFs were also enriched in the lists of TFs predicted to drive the differential expression of genes uniquely altered by rotenone treatment, including CEBPB, STAT3, STAT5A, RELA, SMARCA4 and POU2F2 (Table S10). Coupled to the DMR-DEG TSS analysis, these data suggest that a large fraction of the DEGs had their expression influenced by DNA methylation.

Finally, we evaluated the global relationship between DNA methylation and gene expression. Specifically, we hypothesized that if differential methylation impacts gene transcription, then there is a greater likelihood that DEGs as an unit are differentially methylated at the same time compared to all other genes that are not differentially expressed. To address this, we tested whether gene annotations for DMRs based on their nearest TSS were significantly represented in the overall list of DEGs at different time points. We also tested whether the statistical significance of gene representation was sensitive to discriminating between aging- or exposure-responsive DEGs. What we found was that when evaluating all DEGs, there was significant enrichment for DMRs at the same time relative to the rest of the genome (Fig. 4E, upper panel). Interestingly, the significance seemed to shift to an earlier time point in the rotenone-exposed offspring (Fig. 4E, upper right panel). Indeed, when we segregated the analysis based on ‘aging’ DEGs (Fig. 4E, middle panels) or those uniquely changed in the control or rotenone-exposure (Fig. 4E bottom panels), it became evident that DEGs from the livers of animals that were exposed to rotenone were more significantly differentially methylated at a previous time point. For instance, the methylome at 6 months of age in the treated group resembled that of the 12-month-old animals in the control group (Fig. 4E, right lower panel, see red arrows). These results suggest that the DNA methylation landscape was ‘precociously’ changed by rotenone, likely priming the locus months in advance of the changes in transcriptional output. Similar findings were previously reported for diet-induced obesity in the mouse (23).

### Rotenone alters mitochondrial function in early development and later in life

Since the epigenetic state of the *A^vy^* locus is set prior to the 3-germ layer separation (7, 8, 24), the data on the altered coat-color distribution in the offspring (Fig. 1B) indicate that the first effects of rotenone occurred early in development. As dams were exposed through the diet, it is also expected that rotenone would affect mitochondrial function in various organs of the mother, although likely to different degrees, including the utero and thus the developing embryo. The observed inhibition of oxygen consumption in isolated mitochondria from the livers of mothers exposed to rotenone (Fig. S1B) supports this assumption. In addition, we found increased activity of the mitochondrial superoxide dismutase (MnSOD) in the livers of the treated dams (Fig. 5A). Activation of MnSOD is in line with rotenone inhibiting NADH oxidation at complex I, which is expected to increase superoxide anion production and thus activate the superoxide dismutases (25).

**Figure 5.**
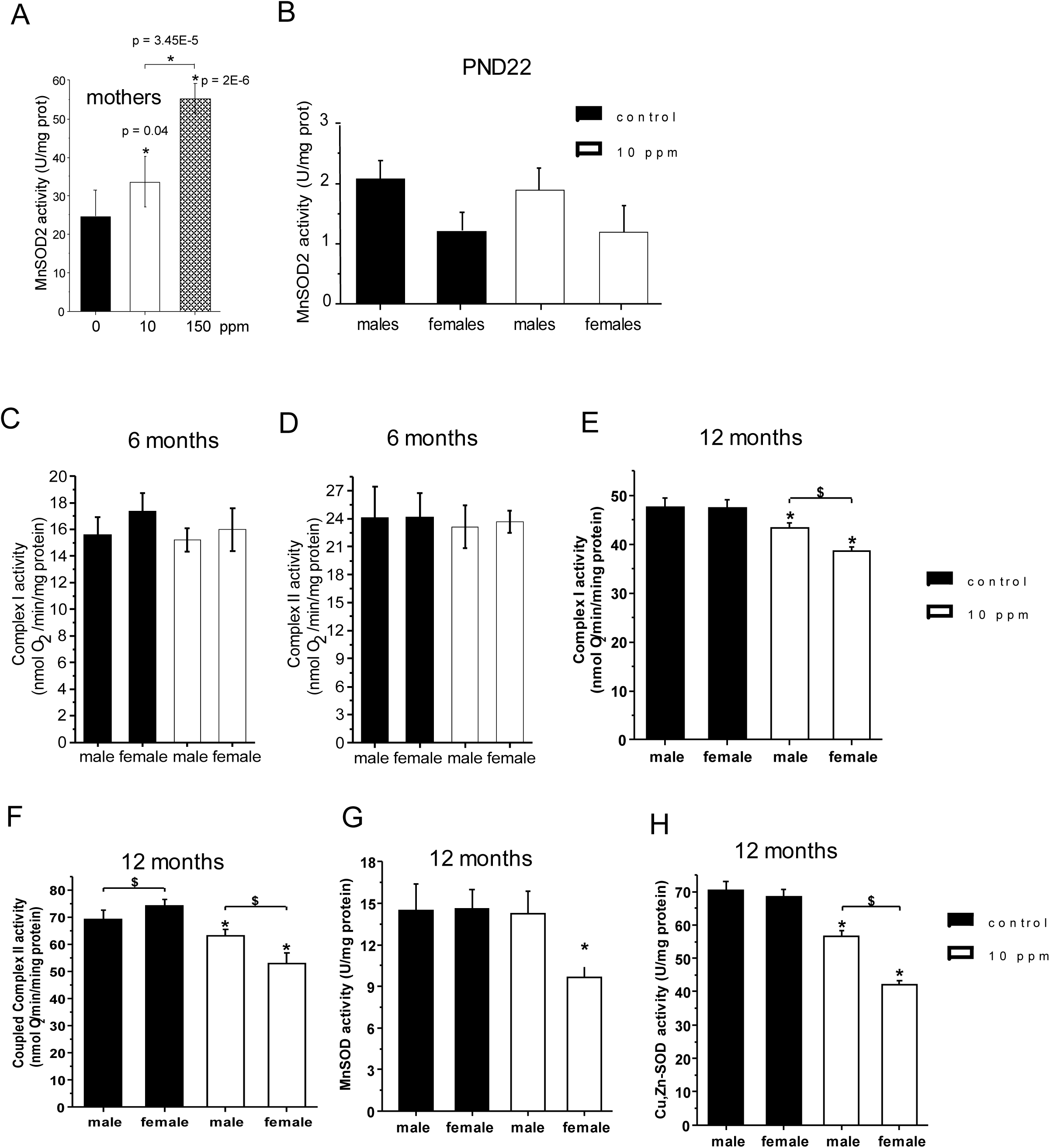
Mitochondrial dysfunction is not present post-natally in the rotenone-exposed offspring until they age. Mitochondrial function was determined in isolated mitochondria from the liver of C57BL/6J mice exposed to control (AIN-93G) or rotenone-containing diets. (A) MnSOD activity was measured spectrofluorometrically in isolated mitochondria from the livers (N=4) of females directly fed rotenone following the inhibition of cytochrome c reduction by a constant flux of superoxide generated by xanthine oxidase. (B) MnSOD activity was gauged as in A in either male and female pups at PND22 N=4. (C-F) Oxygen consumption through complex I (malate + glutamate) or II (succinate + rotenone) was determined in isolated mitochondria from livers using a Clark electrode; N=4. (F) Mn or Cu,Zn-SOD activities were measured in the livers of 12-month-old animals using a spectrofluorometer to follow the inhibition of cytochrome c reduction by a constant flux of superoxide generated by xanthine oxidase. Cyanide was used to selectively inhibit the dismutase activity of Cu,Zn-SOD; N=4/assay/time point. Males and females were assayed separately. All supplemental tables can be accessed through the following hyperlinks: https://orio.niehs.nih.gov/ucscview/Santos/Rotenone/Table S1 - all DMRs.xlsx https://orio.niehs.nih.gov/ucscview/Santos/Rotenone/Table S2 - KEGG DMRs.xlsx https://orio.niehs.nih.gov/ucscview/Santos/Rotenone/Table S3 baseline DEGs.xlsx https://orio.niehs.nih.gov/ucscview/Santos/Rotenone/Table S4 - common aging DEGs.xlsx https://orio.niehs.nih.gov/ucscview/Santos/Rotenone/Table S5 - only controls DEGs.xlsx https://orio.niehs.nih.gov/ucscview/Santos/Rotenone/Table S6 - only rotenone DEGs.xlsx https://orio.niehs.nih.gov/ucscview/Santos/Rotenone/Table S7 - pathology at 12 mo.xlsx https://orio.niehs.nih.gov/ucscview/Santos/Rotenone/Table S8 overlap DEGs vs DMRs at TSS.xlsx https://orio.niehs.nih.gov/ucscview/Santos/Rotenone/Table S9- overlap ENCODE regulatory DNA.xlsx https://orio.niehs.nih.gov/ucscview/Santos/Rotenone/Table S10 - HACER results.xlsx https://orio.niehs.nih.gov/ucscview/Santos/Rotenone/Table S11 - WGBS QC.xlsx

To define the extent to which maternal exposure to rotenone altered mitochondrial function in the pups post-natally, we started by measuring MnSOD activity as surrogate for complex I inhibition at the PND22 time-point; no changes were observed in MnSOD function in either males or females at PND22 (Fig. 5B). Similarly, no changes in MnSOD or complex I activity were found in 3-month-old pups, even when animals were exposed through the mothers to 150 ppm (Fig. S4A and B). Complex I and II activities were not changed in animals evaluated at the 6-month time-point, irrespective of gender, (Fig. 5C an D) but were significantly altered at 12 months (Fig. 5E and F). Activity of MnSOD and the cytosolic Cu,Zn-SOD were also altered at this time point (Fig. 5G and H). These changes were not associated with decreased transcription of genes that make up complex I or complex II or the SODs (Tables S3-6, Fig. S4C), decreased mitochondrial content or of SOD protein levels (Fig. S4D). Interestingly, significant downregulation of genes for metal transporters, including those involved in Zn and Mn that are co-factors of the SODs, were identified in the rotenone-treated cohort (Table S6).

Together, these results lead to three main conclusions: (i) the rotenone-induced mitochondrial dysfunction associated with the DNA methylation alterations observed in the animals up to the age of 6 months was imparted pre-natally, (ii) DNA methylation changes found in 12-month-old animals may have had an additional contribution of mitochondrial dysfunction at the time of the analysis and (iii) that maternal rotenone exposure seems to have facilitated mitochondrial and antioxidant dysfunction in the offspring later in life.

## Discussion

Our data provide unequivocal evidence that perinatal rotenone exposure altered the DNA methylation landscape *in vivo*. This was observed not only at the *A^vy^* locus but across the genome in the liver. Given the effects on the *A^vy^* locus, we assume that the earliest effects of rotenone were in pre-implantation embryos (7, 8, 24). Notably, it is during the pre-implantation stage the first wave of genome de-methylation and re-methylation occurs in the mouse, programming the DNA methylation landscape of the organism (26). Alterations of the epigenome at this early stage of development have been shown to impact the embryo and influence late-life outcomes (27), including in the *A^vy^* model (7, 8, 10). Also interesting is the fact that in the pre-implantation embryo mitochondria undergo dynamic, functional and structural changes, including increasing cristae surface and oxidative phosphorylation activity (27). The main target of rotenone is the site of NADH oxidation at complex I in functional mitochondria.

We assume that rotenone affected DNA methylation by inhibiting mitochondrial respiration, which would be in line with recent literature demonstrating that mitochondrial dysfunction alters this epigenetic mark (1–4). We have not evaluated mitochondrial function in the pre-implantation embryo, when the first effects of rotenone on DNA methylation (*A^vy^* locus) were identified. However, complex I was inhibited in the livers of the exposed dams and it is likely that such effects extended throughout the animal, including in the uterus and thus the embryo. In the pregnant uterus, rotenone could also affect placentation and indirectly the embryo. Nonetheless, litter size and pup weights, parameters known to be affected by placental defects, were not changed in the 10 ppm- exposed cohort (Table 1). Consistent with the effects of rotenone on DNA methylation not being mediated by the placenta, at least in early stages, is the fact that placentation occurs after implantation and hence beyond the developmental window when the effects in the *A^vy^* locus are estimated to occur (7, 8, 24). It is possible that rotenone had off-target effects that drove the epigenetic changes. To date, the only reported non-mitochondrial effects of rotenone involve defects in tubulin polymerization (17), which can disrupt cell division through altered microtubule dynamics and in turn affect organ and animal size - parameters that were not affected in the 10 ppm-treated animals (Table 1). Also, gene expression analysis at PND22 (immediately after rotenone exposure ceased) did not identify changes in the transcription of genes associated with tubulin or its metabolism (Table S3). Finally, to our knowledge, there is no report demonstrating a relationship between tubulin and DNA methylation or the DNA methylation/demethylation machinery, making it unlikely that rotenone exposure altered the epigenome through tubulin defects.

Mechanistically, impaired mitochondrial complex I provided by rotenone could affect DNA methylation through changes in the redox state of the cell, in turn affecting epigenetic modulators that are redox sensitive such as the Ten Eleven Translocation (TET) enzymes (5). Oxidative stress could also modulate DNA methylation by engaging base excision repair (BER), which removes oxidative lesions on the DNA including on methylated cytosines (28). Our data, however, do not seem to support a state of oxidative stress with activation of BER as no signs of oxidative damage were observed in the mothers (Fig. S5 and B), no changes in MnSOD activity were observed at weaning and in 6-month-old animals nor gene expression profiles indicated changes in BER genes. Increased succinate resulting from an inhibited complex I could, alternatively, impair the TETs (29). It is also possible that rotenone by virtue of inhibiting the ETC affected one carbon metabolism and levels of S-adenosyl-methionine, in turn impacting DNA methylation (3). More studies are required to tease out between these possibilities.

That maternal rotenone exposure affected DNA methylation in the offspring that persisted much later in life was clear from our analysis. However, we cannot unequivocally conclude that the DNA methylation changes drove gene expression outputs although our data strongly support a correlation between the two. The relationship between DNA methylation and transcription is complex. For example, the outcomes of DNA methylation for transcription vary depending on whether it occurs on CpG islands, shores or the gene body (30). Also, assigning the ‘correct’ gene to a DMR is still arbitrary, and the TSS of the gene that has the closest proximity to the DMR may not be the one impacted by that differential methylation in the 3D configuration of chromatin within the nucleus. Moreover, epigenetic changes are combinatorial and affect gene expression based on several features including DNA methylation, chromatin accessibility, histone modifications and TF binding (31). Thus, additional experiments such as chromatin configuration capture techniques in the context of rotenone and its removal will be required to establish cause-effect relationships. Despite this, our results demonstrating that effects of developmental exposure to rotenone on DNA methylation and gene expression are still observed later in life suggest that mitochondrial dysfunction has the potential to fundamentally shape long-life health outcomes in unprecedented ways.

In summary, we provide evidence that developmental exposure to rotenone, a well-known environmental pollutant that targets mitochondrial complex I, has effects on the epigenome that are relevant *in vivo*. Given the key role for mitochondria in many different physiological and pathological contexts, including stem cell differentiation, immune cell activation and cancer, these data raise fundamental questions about the broad biological consequences of changes in mitochondrial function to health and disease. This seems particularly relevant in view of the finding that ∼20% of the 10K chemicals evaluated by the NTP impacted mitochondrial function (32, 33) revealing the pervasiveness of environmental mitochondrial toxicants. Our data present a new mechanistic paradigm for which these chemicals can potentially have long-term impacts to human health.

## Online Methods

### *In vivo* animal studies and coat color phenotyping

Female C57BL/6J mice and B6.C3-*A^vy^*/J were purchased from the Jackson Laboratory (Bar Harbor, ME). Animals were housed under controlled and standard conditions of temperature and humidity with a 12 h light/dark cycle. After the first week on regular pelleted mouse chow, NIH-31, animals were switched to the customized AIN-93G diets containing 0, 10 or 150 ppm rotenone *ad libitum*. Customized diets were obtained irradiated from Envigo Inc. (Madison, WI). The B6.C3-*A^vy^*/J male breeders were sacrificed after pregnancy was confirmed while dams were euthanized after weaning. All pups were put on the control diet after PND21 until the end of the experiments (∼18 months). Coat color phenotyping occurred for all animals at PND21, and a fraction of the mice were re-phenotyped at 5- and 12-months. Animals were photographed at the time of initial phenotyping, and the images were independently evaluated and scored for coat color distribution by a total of 5 individuals; final data was reported as the average from the 5 phenotypers. Animals were treated in accordance with the NIH Guide for the Care and Use of Laboratory Animals; the study was revised and approved by the NIEHS Animal Study Proposal review board.

### Analysis of rotenone in the feed

Rotenone was procured from Sigma (lot SLBG7568V, St Louis, MO). The identity was confirmed and the purity (∼ 98%) was determined by mass spectrometry. Prior to final studies, the feasibility of preparing a homogeneous feed formulation and the stability of rotenone in the formulation during storage was established as follows. Rotenone was formulated in AIN-93G meal diet by Teklad/Envigo (Madison, WI). Feed formulation was extracted with acetonitrile and analyzed using a qualified high-performance liquid chromatography (HPLC)-ultraviolet (UV) detection method (linearity >0.999; relative standard deviation (RSD), ≤ 1.8%; relative error (RE), ±≤ 2%; recovery, 103%). Formulation concentration was within 10% of target. In addition, the concentration measured in samples taken from the top, middle, and the bottom of the blender had RSD values of 2.8% demonstrating that the formulation was homogeneous. Aliquots of formulations stored at ambient and refrigerated conditions for up to day 42 were within 10% of day 0 concentration. Similar findings were obtained when simulating the animal room environment, overall demonstrating that the rotenone formulations were stable under the experimental conditions.

### Tissue fractionation and processing

Livers were retrieved surgically from PND22, 6- 12- and 18-month old mice, frozen in dry ice, and stored at −80°C for later processing immediately after sacrifice. Following completion of time course experiment, all specimens were retrieved at once and thawed on ice. Liver tissue from each specimen (50-100 mg each, dry weight) was minced individually with a double-blade on sterile plastic 100-mm plastic dishes while submerged in 5 mL of washing buffer, consisting of chilled 1x PBS supplemented with 0.01% DEPC and 400U of NxGen RNase Inhibitor (Lucigen). Each minced specimen was collected into 15-ml conical polystyrene tubes, vortexed briefly, and spun for 10 min at 1,000×g in a temperature-controlled centrifuge pre-cooled to 4°C. After discarding the supernatant, each sample was rinsed, spun and cleared two additional times in 5 mL washing buffer, then resuspended in 1 mL of washing buffer, transferred to 1.5 mL tubes for fractionation, and pelleted to discard supernatant as described above. Pelleted minced tissue was homogenized further with sterile disposable pestles using a hand-held motor (Kimble) in the presence of 1 mL of Trizol reagent (Invitrogen) and fractionated according to the manufacturer’s instructions to obtain total RNA, genomic DNA, and protein fractions. Yield and purity of nucleic acid fractions were determined by absorbance measurements with a NanoDrop system (Thermo Scientific); protein contents were assessed with colorimetric assays using a commercial Bradford dye-binding method following the manufacturer’s guidelines (Bio-Rad Protein Assay).

### Whole genome bisulfite sequencing (WGBS) library preparation and sequencing

WGBS was performed in liver from animals in the control or rotenone-exposed cohort starting with 1 µg genomic DNA spiked with 50 ng of phage λ, or λDNA (0.5% w/w, Promega), sheared to 300-800 bp size range with a BioRuptor device (Diagenode) for 12-15 cycles (low power setting, 30s on – 90s off) in a total volume of 100 µL inside 0.65-mL clear polystyrene tubes. After confirming shared DNA size range by gel electrophoresis, samples were subjected to bisulfite conversion (EZ DNA methylation kit; Zymo Research) following manufacturer’s instructions. Each sample was ligated and amplified into sequencing libraries with different single-indexed methylated adapters per sample (TruSeq RNA v2, Sets A and B; Illumina) combined with a low-input DNA ligation chemistry (HyperPrep kit, KAPA Biosystems) following the manufacturer’s guidelines. Each individual library was PCR-amplified in technical duplicates for no more than 12 cycles afterwards, and duplicate PCR reaction volumes were collected for DNA purification with size selection by double-sided 0.6×–0.8× SPRI (expected fragment size: 250–450 bp inclusive of sequencing adapters) using AMPureXP magnetic beads (Beckman-Coulter). WGBS library preparation protocol was performed in duplicate, with re-assignment of samples across indices, to screen for batch effect bias and supplement low-quality mappings in bioinformatics post-processing as needed.

### WGBS data processing

General quality control checks were performed with FastQC v0.11.5 (http://www.bioinformatics.babraham.ac.uk/projects/fastqc/). Although some libraries were sequenced in paired-end format, only read 1 was used for consistency across datasets. Raw sequence reads were filtered to retain only those with average base quality score at least 20. Adapter sequence was trimmed from the 3’ end of reads via Cutadapt v1.12 (33) (parameters -a AGATCGGAAGAG -O 5 -q 0 -f fastq). Reads less than 30nt after adapter-trimming were discarded. Filtered and trimmed datasets were aligned to a genome including of the mm10 reference assembly (GRCm38) with the genome sequence of λDNA (NC_001416.1) appended. Mapping was performed via Bismark v0.18.1 (34) at default parameters with Bowtie2 v2.3.0 (35) as the underlying alignment tool. Mapping was first attempted for full length queries (76bp for library A preps, 151bp for library B preps, and 101bp for library C preps). To salvage failed hits due to significant quality concerns, an iterative alignment process was implemented for some datasets. For reads that were not successfully mapped at full length from the library B preps, attempted alignments were done after clipping to 100nt and again after clipping to 50nt. For reads that were not successfully mapped at full length from the library C preps, attempted alignments were done after clipping to 50nt. Mapped hits were deduplicated within libraries. Mapped hits with three or more methylated cytosines in non-CpG context were filtered out since we predicted that they represented incomplete bisulfite conversion. Additional post-alignment read clipping was performed to remove positional methylation bias as determined from QC plots generated with the ‘bismark_methylation_extractor’ tool (Bismark v0.18.1). Specifically, cycle 1 (i.e. 1nt from 5’ end of read) was removed from all mapped hits from library A datasets, and cycles 146-151 (i.e. 6nt from 3’ end of read, unless already clipped prior to alignment) were removed from all mapped hits from library B datasets. Alignments for multiple libraries from the same animal were merged, then hits from mm10 and phage λ were separated. The observed bisulfite conversion rate, which was calculated based on either hits to phage λ or to mouse chrM, averaged ∼99% across all animals in the study. Reads mapped to the mm10 reference genome were used for downstream analysis. All parameters of quality control can be found in Table S11.

### Identification of differentially methylated regions

Differentially methylated regions (DMRs) were identified via MOABS v1.3.4 (36) using merged replicates per sample group. Input data was limited to females with at least two libraries per animal. Pairwise comparisons either between timepoints or between treatment conditions were performed with the ‘mcomp’ function at default parameters. DMRs identified by the M2 method were filtered to require (a) p<0.001, (b) at least 3 CpG sites, and (c) a composite methylation difference of at least 10%, where composite methylation is defined as the total methylated base count at all CpG sites in the defined region divided by the total cytosine base count. A total of 27,562 DMRs in 32 sets were identified from 16 pairwise comparisons of sample groups, with hyper- and hypo-methylated regions in each. For easier assessment of DMR trends, these individual DMR sets were collapsed into groups as follows: aging DMRs in untreated mice (i.e. regions identified as hyper- or hypo-methylated at different ages under 0ppm treatment condition), aging DMRs in rotenone-treated mice (i.e. regions identified as hyper- or hypo-methylated at different ages under 10ppm treatment condition), hypermethylated treatment DMRs (i.e. regions with 10ppm>0ppm methylation level at a given timepoint), and hypomethylated treatment DMRs (i.e. regions with 0ppm<10ppm methylation level at a given timepoint). For each DMR category, DMRs identified by the individual MOABS comparisons and filtered as described above were merged via BEDtools v2.24.0 mergeBed (37) with parameters “-d -2 -o collapse” to generate a set of non-overlapping regions. The two sets of aging DMRs were each further categorized by expression patterns over time. This was accomplished by calculating the composite methylation score for each sample group at each DMR then applying k-means clustering to the row-scaled methylation values. The optimal k for each dataset (k=6 for untreated condition, k=5 for rotenone-treated condition) was determined by the elbow method based on total within-cluster sum of squares tested over k=1 through k=25. One additional cluster was assigned for DMRs with incomplete data; specifically, DMRs with zero cytosine base counts in one or more sample groups. Gene assignments were based on Ensembl transcripts in mouse GENCODE VM18, as downloaded from the UCSC Table Browser (http://genome.ucsc.edu/cgi-bin/hgTables) for mm10 as of December 17, 2018. Each DMR was given a nearest-gene assignment based on distance to TSS, limiting to transcripts from genes with a corresponding Entrez identifier (referenced against ftp://ftp.ncbi.nlm.nih.gov/gene/DATA/GENE_INFO/Mammalia/Mus_musculus.gene_info.gz).

### Motif enrichment analysis using DMRs

Enriched motif analysis was performed with HOMER v4.10.3 (38) for query regions expanded to a minimum width of 200bp centered on the DMR midpoint; these analyses were run using the findMotifsGenome.pl tool with ‘-size given’ (all other parameters default).

### Overlap of DMRs with regulatory DNA

ENCODE data from mouse liver samples was downloaded to evaluate overlap of DMRs with specific genomic marks of interest, including DNase I hypersensitive hotspots (PND0 accession ENCFF054QYT), H3K27ac peaks (PND0 accession ENCFF290MLR and adult accession ENCFF545NYE), H3K4me1 peaks (PND0 accession ENCFF165ZKL and adult accession ENCFF338OJA), H3K4me3 peaks (PND0 accession ENCFF258RQA and adult accession ENCFF420PUU), and H3K9ac peaks (PND0 accession ENCFF587BYL and adult accession ENCFF787EAZ). DNase I hypersensitive hotspots for adult 8-week male liver were downloaded from the UCSC Genome Browser (UCSC Accession wgEncodeEM001720), then converted from mm9 to mm10 coordinates with liftOver from the UCSC utility tools at https://genome.ucsc.edu/util.html. Overlap between query DMRs and peak regions was determined via BEDtools v2.24.0 ‘intersect’. The statistical significance of each overlap was determined by Monte Carlo simulation, where BEDtools v2.24.0 ‘shuffle’ was used to select random regions size-matched to the query DMRs (defined by H3K27ac peaks) followed by overlap assessment via BEDtools v2.24.0 ‘intersect’ for 10,000 iterations. The random region selection excluded chrY, chrM, all non-canonical chromosomes, and any poly-N genomic region longer than 100nt.

### HACER analysis

The list of DMR-derived intervals obtained in mm10 were converted to hg19 using web tool LiftOver at UCSC Genome browser (https://genome.ucsc.edu/). These converted intervals were then used as input for web tool HACER to obtain enhancer annotation (22). The output of HACER, was further processed to generate a summary of TFs and frequency of each TF among these intervals.

### RNA-seq library preparation, sequencing and data analysis

RNA-seq was performed in the livers of females (N=3/group) at PND22, 6-, 12- and 18-month-old animals starting with poly-A-selected RNA from 1 µg of DNAseI-treated total RNA input using a magnetic poly-dT bead hybridization capture (Illumina), eluted in 10µL of nuclease-free H_2_O by heating briefly at 80°C for 2 min, transferred as magnetically isolated supernatant to RNAse-free wells of a 96-well reaction plate, and used directly for rRNA-free library assembly using a KAPA Stranded RNA-Seq with RiboErase commercial kit (KAPA Biosystems) following the manufacturer’s guidelines. Each individual library was PCR-amplified in two technical replicates for no more than 12 cycles afterwards, and duplicate PCR reaction volumes per sample were collected for DNA purification with size selection by double-sided 0.6×–0.8× SPRI (expected fragment size: 250–450 bp inclusive of sequencing adapters) using AMPureXP magnetic beads (Beckman-Coulter).

Libraries were combined in equimolar proportions and sequenced as a single multiplexed library with a NovaSeq 6000 system using an S4 flow cell [Illumina]. Raw sequences were filtered to retain only those with average base quality score at least 20 in both reads of a given read pair. Filtered reads were mapped to the mm10 reference assembly (GRCm38) via TopHat v2.0.4 with parameters ‘--b2-sensitive --library-type fr-unstranded -g 1 --mate-inner-dist -40 --mate-std-dev 50’ (39). Resulting hits were coordinate-sorted and deduplicated with Picard tool suite v1.96, then talliled for further data analysis with SeqMonk, version 37.1 (Andrews, S. SeqMonk, 2007: http://www.bioinformatics.babraham.ac.uk). Composite per kilobase per million reads (RPKM) counts within genomic coordinates of genes were used to calculate gene expression differences.

Detection of differentially expressed genes (DEG) across time×treatment groupings was performed using weighed two-way ANOVA (gene×grouping) based on log2-transformed fold-change (log2FC) measurements relative to gene RPKM grand-means over all specimens; N=24 (3 biological replicates per time×treatment grouping). Gene-wise log2FC values were weighed by a relative metric of sequencing representation (cumulative hazard of significance scores from gene-wise RPKM rate modeling with an exponential distribution and inverse link function) as previously described (40). Genes were retained for post hoc pairwise analysis when significance level *p*<0.05 after multiple comparison adjustment (41), then filtered against a minimum gene-wise effect size δ_log2FC_>0.3×σ_log2FC_ and *post hoc* pairwise significance (Student’s t-test *p*<0.05) for log2FC differences at each timepoint against PND22 among specimens in the same treatment group, as well as log2FC differences at PND22 between treatment groups. For gene-level effect size filtering, δ_log2FC_=0.3×σ_SSR_ is 5% of the 6σ-spread log_2_FC regression error with respect to a gene’s grand mean [where (σ_SSR_)^2^=(SSR_log2FC_)/(N-1)] compared to 5% of the 6σ-spread in measurement error about the mean log2FC of time×treatment groupings in the gene [where (σ_log2FC_)^2^=(SSE_log2FC_)/(N-1)].

### Mitochondrial oxygen consumption measurements

Mitochondria were isolated by differential centrifugation (42) from freshly harvested livers finely chopped into ice-cold mitochondria isolation buffer (10 mM HEPES, pH 7.4, 0.25 M sucrose, 1 mM EGTA, 0.5% BSA) using a Dounce homogenizer. Oxygen consumption was monitored using a Clark electrode and substrates for complex I (5 mM malate + 5 mM glutamate) or II (5 mM succinate + 20 µM rotenone)-driven respiration; basal mitochondrial respiration was recorded (0.5 mg/mL mitochondria) for 1-2 min prior to the addition of the substrates; ADP (200 µM) and CCCP (20 µM) were also added to determine state 4/3 and maximal respiration, respectively. Each analysis was done with N=4 per time point.

### Superoxide Dismutase Activity

Superoxide dismutase activity was measured in isolated mitochondria through the inhibition of cytochrome c reduction by a constant flux of superoxide radicals generated by xanthine oxidase in the presence of xanthine (43). In whole tissue homogenates, after total SOD activity was determined using the same protocol as above, cyanide was used to selectively inhibit Cu,Zn-SOD and define MnSOD activity only; these values were subtracted from the total to determine Cu,Zn-SOD contribution.

## Data Availability

The WGBS and RNA-seq data described in this manuscript have been submitted to the NCBI Gene Expression Omnibus (GEO) under accession GSE136417.

## Acknowledgments

We would like to acknowledge the support of all members of the Animal Care Group and staff from the Genomics Core both at NIEHS and at NISC. We thank RTI International for their work on the chemical analysis under contract HHSN273201400022C, and Lorenzo Trovero, Valentina Trovero and Ashlyn Jacobs for their help in scoring the coat color of the animals using the photographs. We would also like to acknowledge Dr. Douglas Ganini da Silva for technical assistance in the mitochondria functional assessments and Dr. Grace Kissling for the power analysis that defined the number of animals needed for the coat color analysis. Finally, we are thankful to Drs. Paul Wade and Carmen Williams for critical reading of the manuscript. This research was supported by the NIH, National Institute of Environmental Health Sciences, Division of Intramural Research.

## Author Contributions

OAL and FX performed all experiments, DG provided technical support and performed Western blots. TW and SG analyzed the genomics data, including performing HOMER, HACER and ENCODE overlaps. VG and SW coordinated the analytical chemistry of rotenone concentrations and stability in the diet. RPW and JHS conceptualized the study, oversaw the implementation of the experiments and the analysis of data. JHS was the lead writer of the manuscript.

## Competing Interests

The authors declare no competing interests.

## Figure Legends Supplemental Figures

**Figure S1.**
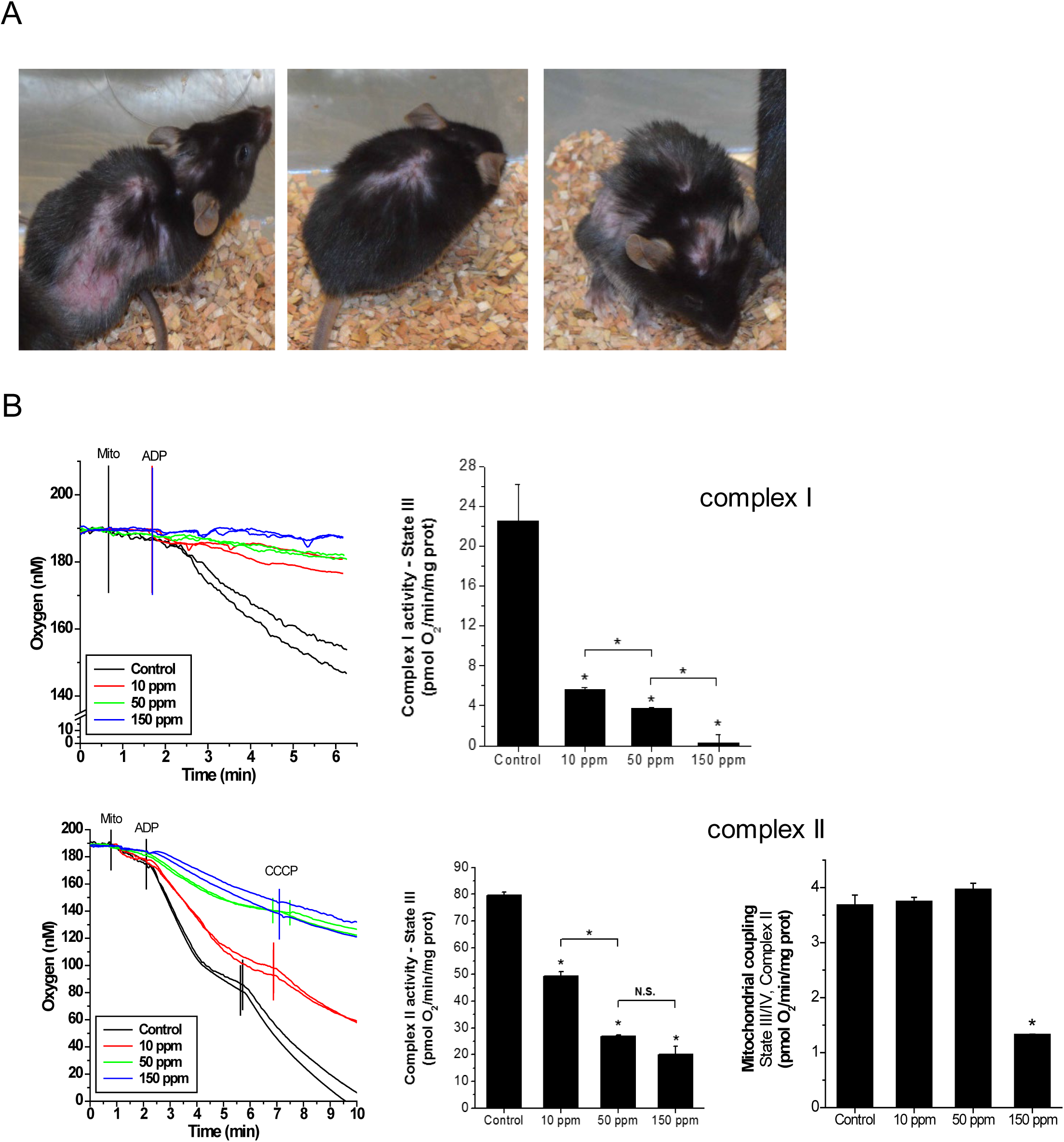
Effects of rotenone exposure. (A) Representative images of detrimental effects of rotenone in the cohort directly exposed to 600 ppm through the diet for 2 days; 3 different animals are depicted. (B) Oxygen consumption in isolated mitochondria from the liver of female C57BL/6J mice that were fed AIN-93G (control) or AIN-93G containing rotenone at the concentration of 10, 50, or 150 ppm. N=4 per group. Polarographic traces of oxygen consumption through complex I (malate + glutamate; upper left panel), which were used to calculate the maximal complex I activity (State III, right upper graph). Lower left panel shows polarographic traces of oxygen consumption through Complex II (succinate + rotenone), which was then used to calculate maximal complex II activity (State III) and mitochondrial coupling, middle and right graphs, respectively. Coupled mitochondrial respiration was calculated based on CCCP measurements.

**Figure S2.**
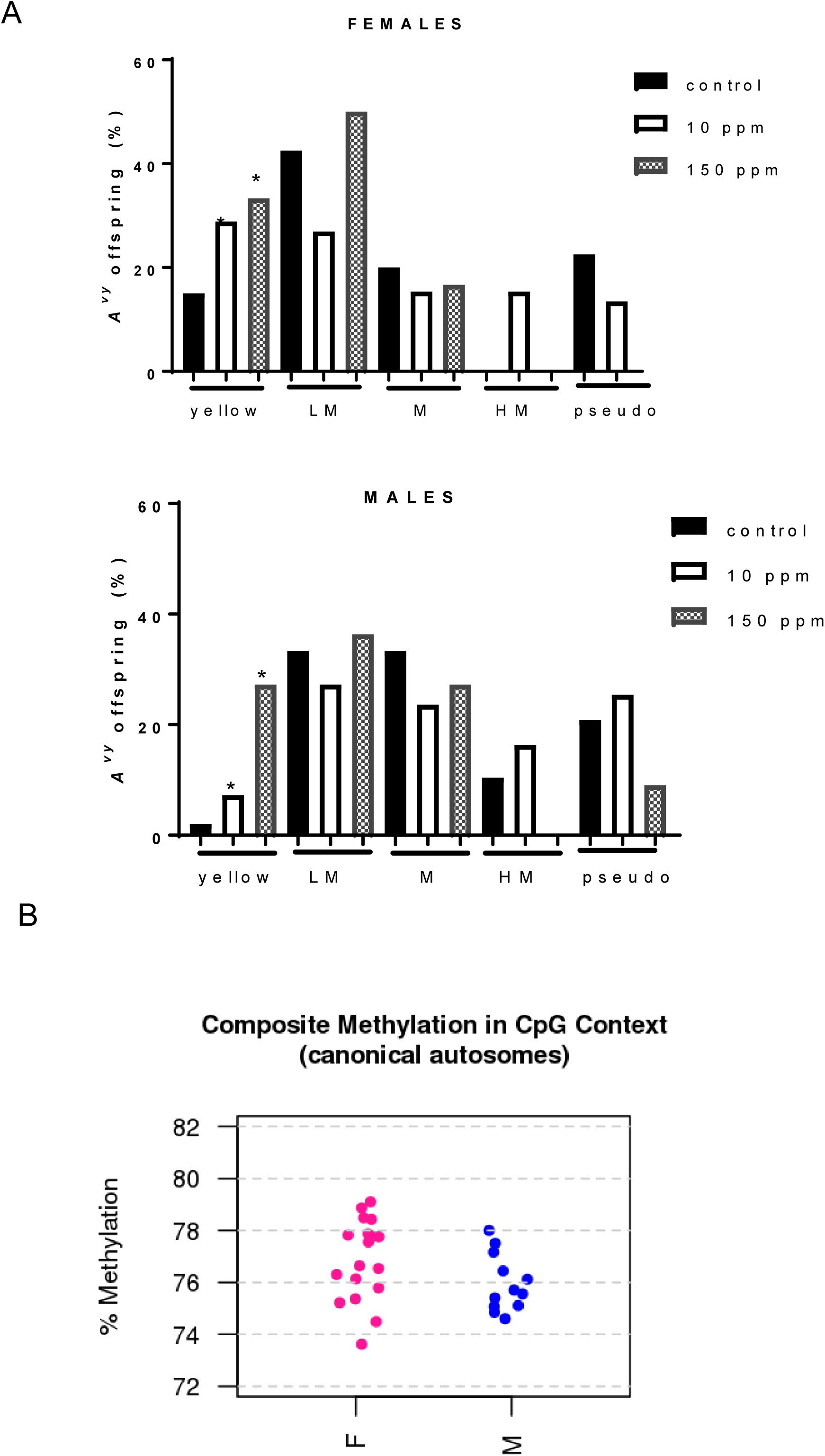
Maternal rotenone exposure alters the frequency of yellow *A^vy^* offspring in both females or males. (A) The frequency of animals in each of the 5 categories of the *A^vy^* coat color phenotype was calculated based on the number of female or male animals in that category relative to the total number of mutant females (N=98) or males in (N=114) the cohort in each experimental group. Y = yellow, LM – lightly mottled, M – mottled, HM – heavily mottled; pseudo – pseudoagouti. Chi-square test was applied to determine significance of coat color distribution differences across doses *p=0.023. (B) Composite global methylation in CpG context using a total of 18 females or 12 males per group at PND22 and at 6-month-old animals. Methylation is defined as total methylated C bases divided by total mapped C bases; depicted are aggregated scores calculated for each sample (shown as dots). Pink = females; blue = males. Only autosomes were evaluated.

**Figure S3.**
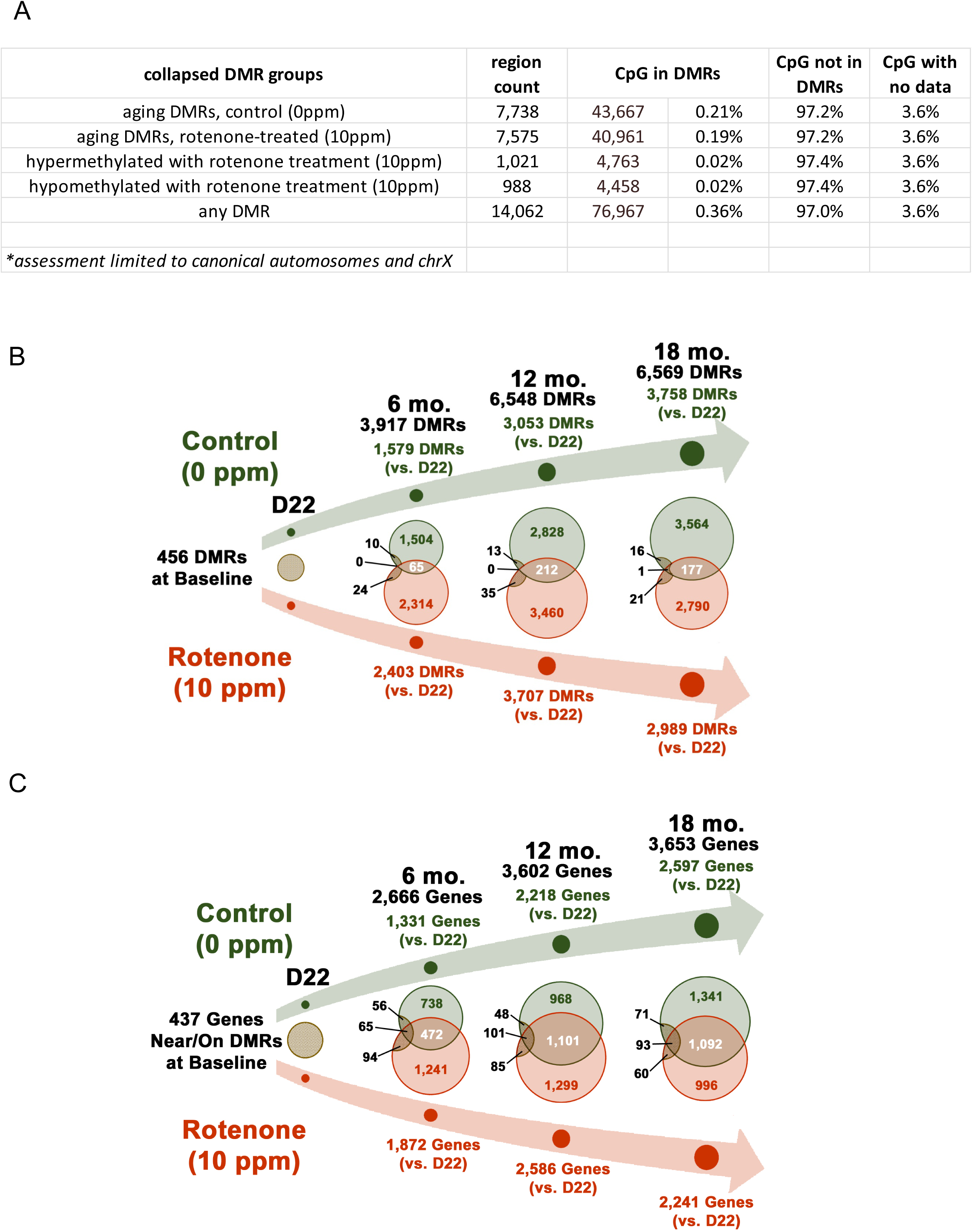
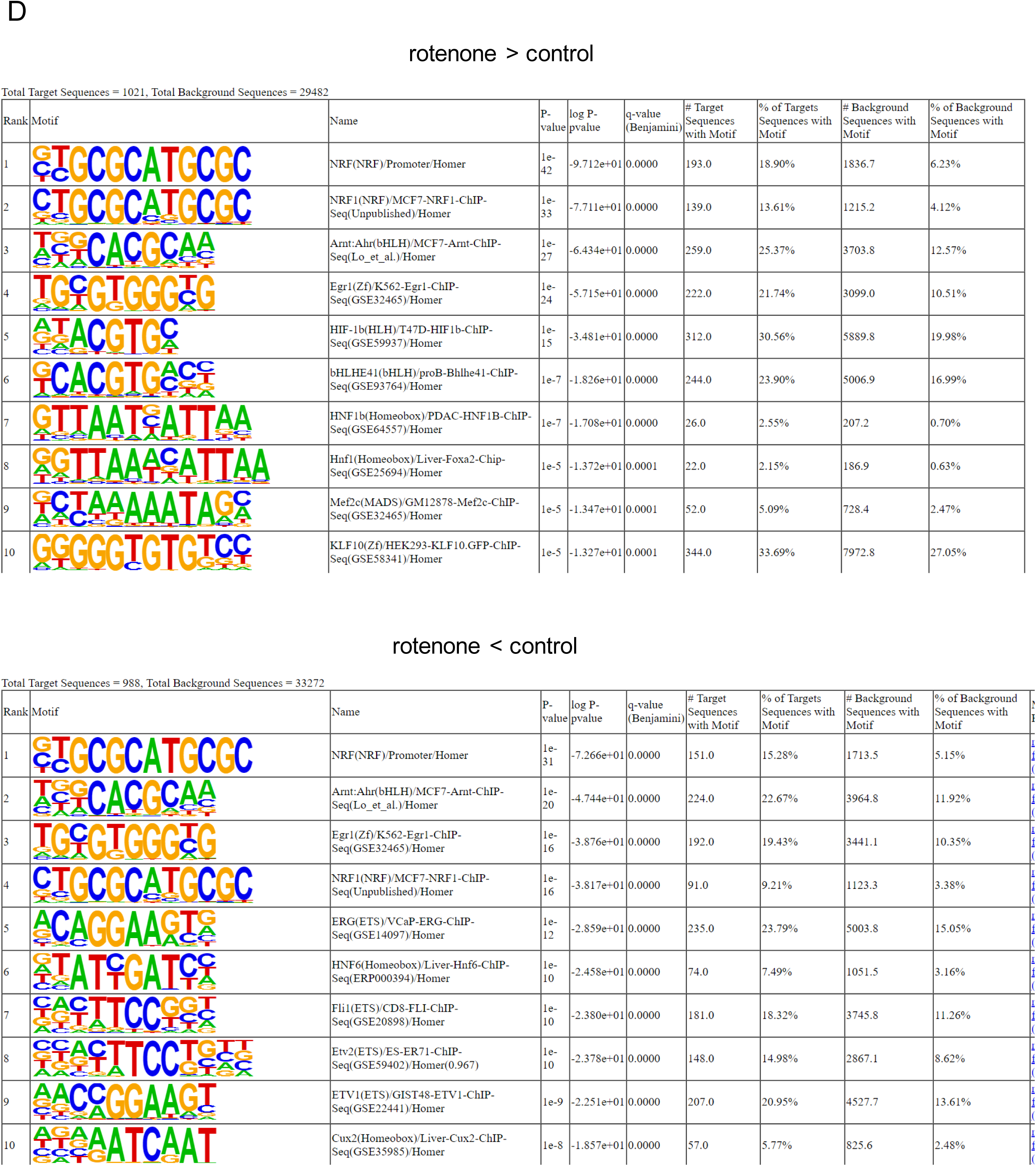
Developmental rotenone exposure modulates the aging liver DNA methylome. (A) The percentage of the genome represented as differentially methylated region (DMR) was calculated based on the full set of CpG sites (limited to only chr1-19,X) in all samples relative to having no data in any of the samples, data but not in a DMR or data in a DMR. (B) Venn diagram depicts the overlap of DMRs at each time point between experimental groups. Black numbers on top depict all DMRs identified. Baseline DMRs were obtained by comparing rotenone to control cohort at post-natal day 22 (D22). Green depict the number of DMRs identified in the control cohort and in orange those identified in the rotenone-fed counterparts. Data at each time point are relative to D22 within each group. The black circles show the number of DEGs identified at baseline that were still differentially expressed at later times and the amount that was common to both groups; number above or below these circles (in black) depict baseline DMRs unique to control (upper) or rotenone-fed cohort (lower). (C) Same as B but with overlap of DMRs based on the nearest TSS of genes. (D) Top 10 enriched transcription factor (TF) motif in rotenone hyper (upper) or hypomethylated (lower) DMRs relative to control according to HOMERv4.9.1.

**Figure S4.**
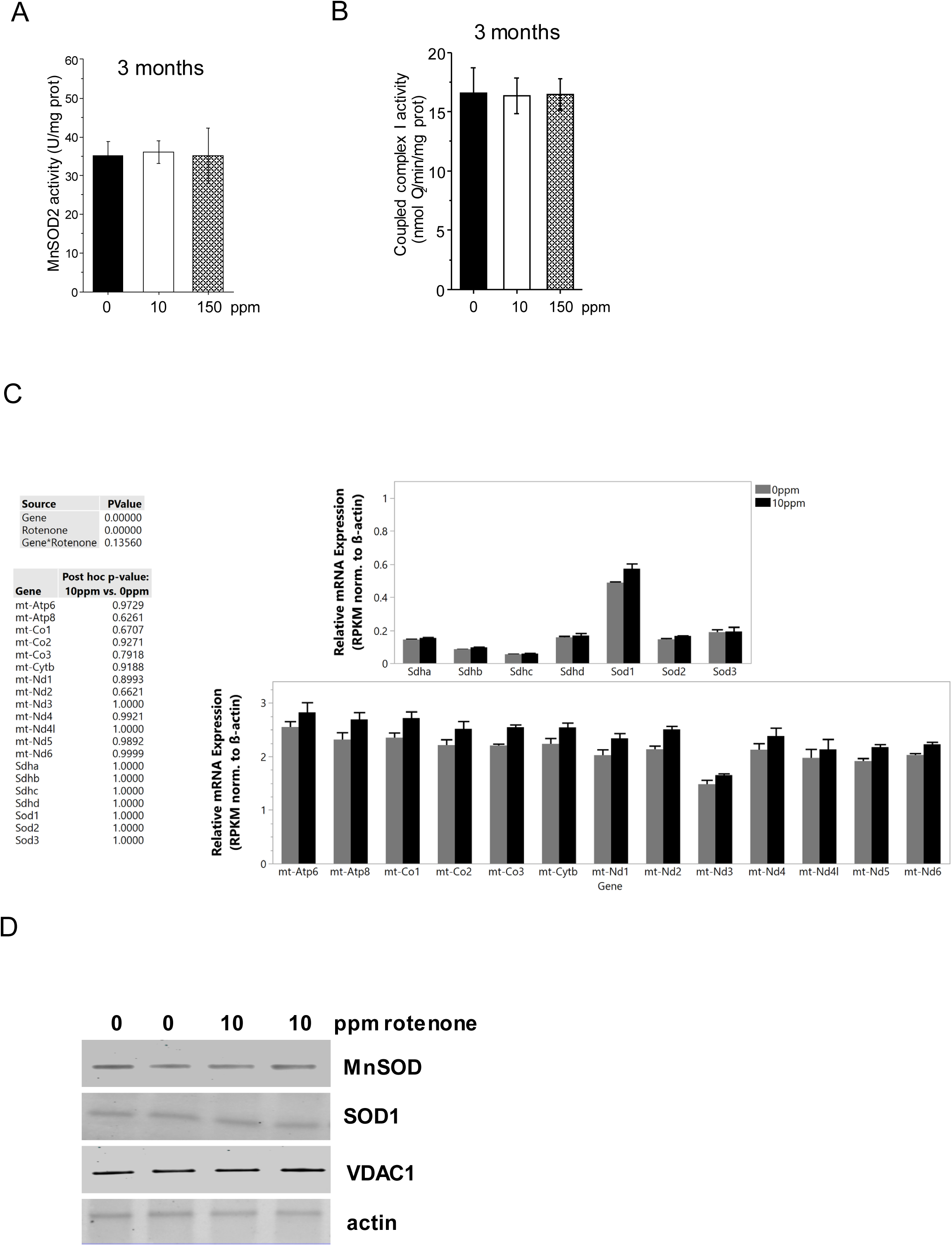
Perinatal rotenone exposure does not alter mitochondrial or SOD content. (A) MnSOD activity was measured spectrofluorometrically in isolated mitochondria from the livers of animals exposed to control, 10 or 150 ppm rotenone-containing diets, (N=4). (B) Oxygen consumption through complex I (malate + glutamate) was determined in isolated mitochondria from livers of animals exposed to the different diets using a Clark electrode; N=4. (C) RNA-seq counts for actin, succinate dehydrogenase, SODs and mtDNA-encoded genes at 12-month-old animals. Each gene was normalized to actin; grey bars = controls, black bars = rotenone. (D) Whole tissue homogenates were obtained from independent animals at 12 months and the SODs and VDAC1 (a mitochondrial membrane protein) were probed by Western blotting; acting was used as loading control and VDAC1 to inform changes in mitochondrial content.

**Figure S5.**
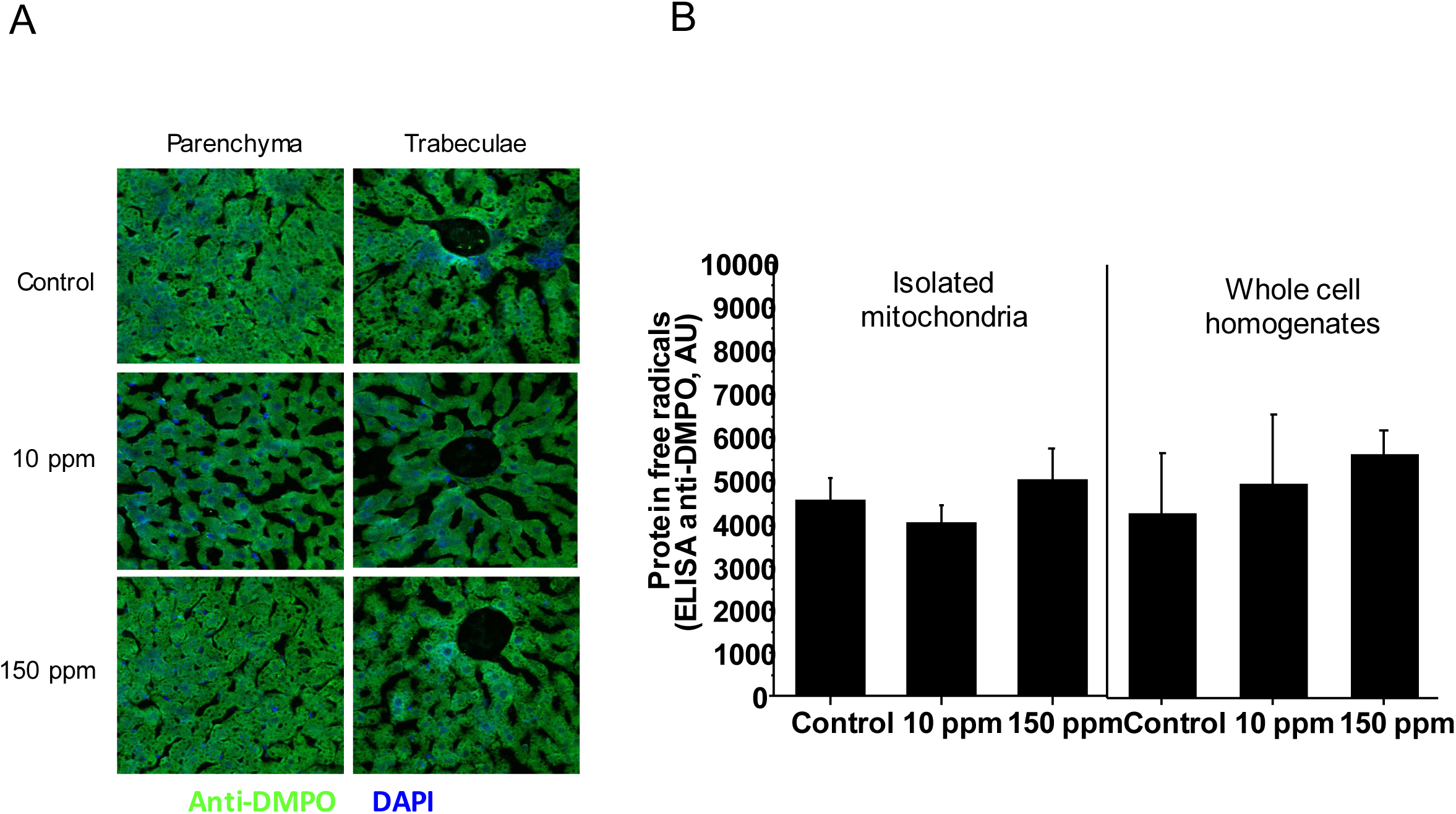
Rotenone exposure does not lead to signs of oxidative damage in the livers of directly exposed damns. (A) Animals were injected with the spin trapper DMPO (5,5-Dimethyl-1-pyrroline N-oxide) 2h prior to sacrifice. Livers were collected from control females or those directly treated with rotenone and an antibody against DMPO was used to detect free-radical intermediates *in situ*. Data are representative of N=3. (B) ELISA was performed to detect protein free radicals using the anti-DMPO antibody in isolated mitochondria or whole tissue lysates from livers of control or rotenone-exposed dams; N=3 per dose.

