## Appendix for "Maternal exposure to a mitochondrial toxicant results in life-long alterations in DNA methylation and gene expression in the offspring"

16-017B1:

A: Liver- Congestion, focally extensive, mild
B: Kidney -There are no significant findings (NSF).
C: Heart- There are several scattered, small areas of cardiomyocyte mineralization.
D: Lung -NSF.
E: Skin - There are multifocal areas of mild to moderate epidermal acanthosis, some of which are subjacent to serocellular crusts. There are occasional foci of full thickness epidermal ulceration, with densely packed neutrophils admixed with fibrin and debris between the dermis and the overlying crust. Adjacent to these crusting and/or ulcerated areas, there is multifocal, compact orthokeratotic hyperkeratosis extending into follicles. Occasional apoptotic keratinocytes are present in both the basilar and middle regions of the epidermal and follicular epithelium. Hair shafts and follicles are rarely present in complete long section, but when they are, they are frequently broken at the follicular ostia. There is mild to moderate superficial dermal edema. An interstitial inflammatory cell infiltrate consisting predominantly of lymphocytes and plasma cells with occasional mast cells is present in all layers but concentrated towards the deeper regions of the dermis. This inflammatory population transitions towards medium to large numbers of perivascular to interstitial neutrophils and fewer histiocytes as dermis transitions towards subcutis. There are occasional foci of loose, globular, light brown pigment (suspect melanin, pigmentary incontinence).

**Diagnosis**: Haired skin- Dermatitis, crusting and ulcerative, lymphoplasmacytic, multifocal with orthokeratotic hyperkeratosis, acanthosis, furunculosis and dermal edema, subacute to chronic, moderate

16-017B2
A: Liver -There is a locally extensive, irregularly shaped region with hepatocytes exhibiting moderately increased amounts of intracellular glycogen and lipid. Congestion, focally extensive, moderate
B: Kidney- NSF.
C: Heart- NSF.
D: Lung- Sinus histiocytosis, moderate, mediastinal lymph nodes
E: Skin- Acanthotic epidermal hyperplasia, focal, with orthokeratotic hyperkeratosis, subacute to chronic, mild

16-017B3
A: Liver- NSF.
B: Kidney - NSF.
C: Heart, lymph node- Sinus histiocytosis, mild, mediastinal lymph nodes
D: Lung - Congestion and atelectasis, multifocal, mild to moderate
E: Skin- In one section, there is a focal area of moderate acanthotic epidermal hyperplasia subjacent to a serocellular crust, with adjacent areas exhibiting mild orthokeratotic hyperkeratosis. There is occasional hydropic degeneration of keratinocytes. There is a perivascular to interstitial inflammatory infiltrate concentrated towards the deep dermis consisting predominantly of neutrophils with fewer

lymphocytes, plasma cells, and histiocytes. Adnexa are not present in the area subjacent to the crust, and there are occasional loose foci of presumptive melanin.

In this same section, there is another area with full thickness epidermal ulceration and partial to full thickness dermal necrosis, with deeply eosinophilic, coalescing collagen fibers admixed with streaming nuclear and cytoplasmic debris. This area is generally devoid of adnexa, with only a few remnants of follicular bulbs remaining, one of which contains presumptive loose melanin pigment. Adjacent to this area, there is a small serocellular crust, and alongside the other margin, there is mild acanthotic hyperkeratosis and compact orthokeratotic hyperkeratosis

Other sections are generally devoid of significant findings, characterized only by occasional minimal epidermal hyperplasia and/or mild superficial dermal edema.

**Diagnosis**: Dermatitis, crusting, ulcerative, & necrotizing, multifocal & perivascular, neutrophilic, with acanthosis & compact orthokeratotic hyperkeratosis, subacute to chronic, moderate

16-017B4
A: Liver- NSF.
B: Kidney - NSF.
C: Spleen and Pancreas- - NSF.
D: Lung - NSF.
Mediastinal lymph node- A mediastinal lymph node is expanded and effaced by a densely cellular, poorly demarcated, multinodular mass. Sheets of round cells supported by sparse fibrovascular stroma have distinct borders and scant to low amounts of granular, eosinophilic cytoplasm. Their round nuclei have variably distinct, sometimes multiple nucleoli with coarsely stippled chromatin. Approximately 2-3 mitoses are present per 400x field. Anisokaryosis and anisocytosis are mild to moderate, and there are intermediate numbers of apoptotic cells.

**Diagnosis**: Lymphoma, mediastinal lymph node

E: Skin- Two sections of haired skin are present. In one section, there is regional, mild to moderate acanthotic epidermal hyperplasia associated with generally basket weave but occasionally compact mild to moderate orthokeratotic hyperkeratosis. In these areas, there is mild, multifocal hydropic degeneration, predominantly in the stratum basale. In the stratum granulosum, there is moderate, variable keratinocyte hypertrophy, as well as rare apoptotic keratinocytes. There are multifocal regions of mature fibroblasts abutting the overlying hyperplastic epidermis and occupying the Grenz zone (fibrosis). Adnexal units are generally reduced in these areas though are occasionally entrapped by the fibrous connective tissue. Occasionally the hair follicles exhibit moderate hyperkeratosis. In regions adjacent to the fibrosis, there is a predominantly histiocytic interstitial inflammatory cell infiltrate that occasionally forms a lichenoid pattern, obscuring the dermal-epidermal junction. There is mild generalized superficial dermal edema, with low numbers of scattered interstitial lymphocytes and plasma cells.

The other section exhibits several multifocal areas with similar though generally less severe findings. There is a locally extensive area however, with marked epidermal acanthotic hyperplasia, marked predominantly compact though occasionally basket weave, orthokeratotic hyperkeratosis extending downward into and also expanding follicles, and deep, irregular rete pegs that contain cells with increased mitotic figures (regeneration). There are regions of variable (mild to moderate) fibrosis in the Grenz zone. Medium to large numbers of histiocytes, with fewer lymphocytes and plasma cells are scattered throughout the underlying dermis, which also is characterized by increased collagen density.

**Diagnosis**: Haired skin -Dermatitis, histiocytic & lymphoplasmacytic, multifocal & interstitial, with dermal fibrosis, epidermal acanthotic hyperplasia, & orthokeratotic hyperkeratosis, chronic, moderate to marked

16-017B5
A: Liver - NSF.
B: Kidney - NSF.
C: Spleen- NSF.
D: Lung -Atelectasis & congestion, multifocal to coalescing, moderate
E: Skin- There is a focally extensive area of serocellular crusting dermatitis overlying moderate epidermal acanthotic hyperplasia. Deep to this area, there is an interstitial inflammatory cell aggregate concentrated towards the deep dermis though present in all areas consisting predominantly of neutrophils with fewer lymphocytes and plasma cells. Within and adjacent to these areas, expanded hair follicles contain clear space (edema) and fibrillar eosinophilic material (fibrin). Adjacent to the crusting dermatitis, there is moderate compact orthokeratotic and parakeratotic (regionally dependent) hyperkeratosis that extends into hair follicles. Deep to these areas, in the subcuticular adipose tissue, there are moderate numbers of perivascular to interstitial neutrophils and histiocytes, with fewer lymphocytes and plasma cells, along with coalescing areas of basophilic granular material (fat necrosis). Other areas of the tissue are occasionally affected by mild epidermal acanthotic hyperplasia and mild orthokeratotic hyperkeratosis.

**Diagnosis**: Haired skin -Dermatitis, interstitial & multifocal, neutrophilic, with acanthotic hyperplasia, orthokeratotic & parakeratotic hyperkeratosis, & locally extensive neutrophilic & histiocytic steatitis, moderate, chronic-active, moderate

16-017B6
A: Liver - NSF.
B: Kidney- There are sporadic areas with mild tubular ectasia and intraductular cytoplasmic blebbing.
Adrenal Gland- NSF.
C: Spleen- Congestion, multifocal to coalescing, mild
D: Lung- Atlectasis & congestion, locally extensive, moderate

The most significant and consistent finding was an ulcerative and crusting dermatitis ranging from acute to chronic (& healing) lesions. No obvious evidence of an infectious agent was detected in these sections, though histology is an insensitive method to detect such agents as compared to culture or PCR. Special stains screening for dermatophytes and bacteria did not reveal any evidence of such etiologies.

Depending on distribution of the lesions on the mice, and if the mice are pruritic, these lesions may be consistent with C57BL ulcerative dermatitis syndrome. This syndrome, for which C57BL mice are at increased risk, has an unknown etiology, though environmental factors such as temperature, diet (increased fat and calories), and humidity are thought to play a factor. It is characterized by pruritic self-trauma leading to ulceration that can progress to epidermal crusts, dermal necrosis and fibrosis leading to potentially debilitating dermal contractures that limit mobility. Most commonly these lesions occur on the head and dorsal thorax. In more severely affected cases, there can be associated lymphadenopathy and splenomegaly. We cannot say with certainty, though we feel it to be doubtful that the nodal lymphoma diagnosed in one mouse was related to potential (we did not have a slide of skin for this mouse) dermal changes (given the lack of other nodal and splenic changes). As opposed to fighting wounds and primary skin infections, these lesions tend to respond poorly to treatment, though a recent publication suggests 0.005% sodium hypochlorite may be more successful than other treatments. A system to score ulcerative dermatitis in live mice is available.

As an aside, when you submit sections of skin for histopathologic evaluation, please section in the plane of hair growth. This allows us to visualize the entire hair follicle in cross section and make a more complete assessment. We are happy to assist you with this if so desired.

Our Pubmed literature search did not find evidence of any reported link between rotenone and dermal lesions.
We consulted with the dermatopathologist at North Carolina State University's College of Veterinary Medicine on this case.

Potentially Useful References:
Hampton AL et al. Progression of ulcerative dermatitis lesions in C57BL/6Crl mice and the development of a scoring system for dermatitis lesions. *Journal of the American Association for Laboratory Animal Science*. 51(5); 586-593, 2012.
Kastenmayer RJ, Fain MA, Perdue KA. A retrospective study of idiopathic ulcerative dermatitis in mice with a C57BL/6 background. *Journal of the American Association for Laboratory Animal Science*. 45(6); 8-12, 2006.
Michaud CR et al. Comparison of 3 topical treatments against ulcerative dermatitis in mice with a C57BL/6 background. *Comparative Medicine*. 66(2); 100-104, 2016.
Neuhaus B et al. Experimental analysis of risk factors for ulcerative dermatitis in mice. *Experimental Dermatology*. 21; 710-720, 2012.
Sargent JL, Koewler NJ, Diggs HE. Systematic literature review of risk factors and treatment for ulcerative dermatitis in C57BL/6 mice. *Comparative Medicine*. 65(6); 465-

472, 2016.
